# Age differences in electrocortical dynamics during uneven terrain walking

**DOI:** 10.1101/2025.10.13.682080

**Authors:** Chang Liu, Erika M. Pliner, Jacob Salminen, Ryan J. Downey, Jungyun Hwang, Arkaprava Roy, Ryland Swearinger, Natalie Richer, Chris J. Hass, David J. Clark, Todd M. Manini, Yenisel Cruz-Almeida, Rachael D. Seidler, Daniel P. Ferris

## Abstract

Walking on uneven terrain becomes more difficult as we age, and gait becomes less automatic. Using mobile brain imaging via high-density electroencephalography (EEG) can provide insight into the neural mechanisms contributing to reduced mobility capability with aging. The objective of this study was to quantify age differences in electrocortical dynamics during uneven terrain walking, both averaged across many strides and variations within a stride. We included 31 young adults and 71 older adults for analysis. All participants walked on an uneven terrain treadmill with four levels of terrain difficulty at their self-selected speed. Compared to younger adults, older adults exhibited a greater increase in step duration variability and mediolateral sacral excursion variability as the terrain became more uneven. We identified multiple brain regions involved during walking on uneven terrain. Regardless of age group, walking on uneven terrain compared to flat terrain led to a widespread change of electrocortical dynamics in the brain, especially in the alpha (8–13Hz) and beta (13–30Hz) band power. In the parieto-occipital region, younger adults experienced a greater reduction in alpha and beta power with increasing terrain unevenness compared to older adults. We also assessed how intra-stride power fluctuations changed with terrain unevenness and age group. Greater intra-stride power spectral fluctuations in the occipital area were associated with greater terrain unevenness for younger adults, but not for older adults. In summary, older adults showed a greater increase in gait variability than younger adults as the terrain became more uneven, but exhibited a lack of modulation of parieto-occipital activity in response to terrain unevenness. The lack of task-related power modulation may suggest reduced cortical network flexibility in older adults. The absence of increased parieto-occipital activity when walking on uneven versus flat surfaces in older adults may also indicate that, unlike younger adults, older adults already heavily rely on visual processes during flat surface walking and may therefore have reduced occipital modulation range remaining to cope with the visuomotor processing demands of walking on uneven surfaces.

## Introduction

One-third of older adults in the United States report mobility limitations (Musich et al., 2018), which is estimated to incur more than $42 billion in healthcare costs (Hardy et al., 2011). Walking ability is a vital indicator for assessing mobility limitations (Brown & Flood, 2013; Groessl et al., 2007). Older adults walk more slowly, have reduced endurance, exhibit altered kinetics and joint kinematics, and are at a greater fall risk than younger adults (Akyol, 2007; Boyer et al., 2023; Frimenko et al., 2015). When walking in daily life, the ability to walk safely on uneven terrain, such as from grass to a concrete sidewalk, is essential for community-dwelling older adults. Walking on uneven surfaces poses a greater risk of falls for older adults compared to walking on flat ground. Uneven terrain causes older adults to walk slower, take shorter steps, and increase their movement variability (Downey et al., 2022; Marigold & Patla, 2008; Thies et al., 2005). These mobility changes may lead to avoidance of community ambulation and diminish quality of life, highlighting the importance of understanding the mechanisms contributing to uneven terrain mobility limitations with aging.

The underlying neural mechanisms contributing to age differences in uneven terrain walking are not well understood. The challenge of walking on uneven terrain increases with age, as walking on flat ground is already less automatic and relies more on cognitive control for older adults (Clark, 2015; Shah et al., 2023). Prior studies using functional near infrared spectroscopy (fNIRS) have reported overaction of the prefrontal cortex in older adults during walking, which suggests a need for greater cognitive control (Belli et al., 2021; Clark, 2015; Clark et al., 2014; Holtzer et al., 2011). It remains unclear how aging impacts activity in other brain regions, although we know that deterioration in motor and sensory systems, visuospatial processing, and sensorimotor integration impacts walking function for older adults (Franz et al., 2015; Sato & Choi, 2022; Seidler et al., 2010).

Innovations in brain imaging using high-density electroencephalography (EEG) allow the direct yet non-invasive assessment of brain activity with high temporal resolution (in milliseconds) during walking and other behaviors (Gwin et al., 2011; Richer et al., 2024; Yokoyama et al., 2021). Each of the EEG spectral power bands (theta, alpha, beta) provides unique insight into brain function during human locomotor tasks (reviewed by Richer et al., 2024). Increasing demand for balance control is consistently associated with an increase in theta power (3 - 8 Hz) and decreases in alpha (8 – 13 Hz) and beta (13 – 30 Hz) power (Blum et al., 2022; Bruijn et al., 2009; Liu et al., 2024; Sipp et al., 2013). Additionally, intra-stride EEG power fluctuations are thought to reflect sensorimotor processing or integration during gait (Guo et al., 2024; Seeber et al., 2014). Intra-stride EEG power fluctuations localized to the sensorimotor cortex have exhibited event-related synchronization during the double support phase and desynchronization during the swing or single support phase of gait (Artoni et al., 2017; Bradford et al., 2016; Bruijn et al., 2015; N. A. Jacobsen & Ferris, 2023; Liu et al., 2024; Zhao et al., 2022). Such power fluctuations became more pronounced when participants were asked to step on precise targets when walking on a treadmill, likely due to increased visuomotor processing for controlling foot placement (Oliveira et al., 2018; Yokoyama et al., 2021). Taken together, quantifying both electrocortical changes at different bands and the intra-stride power spectral fluctuations could enable a comprehensive profile of the influence of older age on brain dynamics during uneven terrain walking.

The objective of this study was to quantify age differences in electrocortical dynamics during uneven terrain walking. We used a high-density EEG system to quantify the gait-related brain activity when younger and older adults walked on uneven treadmill terrains with four levels of difficulty at a preferred walking speed. Our hypotheses were based upon our previous finding that as terrain unevenness increases, younger adults demonstrated a decrease in alpha and beta spectral power in the sensorimotor and posterior parietal areas and an increase in theta power in the mid/posterior cingulate area (Liu et al., 2024). Here, we hypothesized that older adults would exhibit a greater increase in spectral power with terrain unevenness compared to younger adults in theta bands (decrease for alpha and beta) as older adults tend to recruit more neural resources during gait and balance (Hawkins et al., 2018; Holtzer et al., 2011). We also hypothesized that older adults would demonstrate greater intra-stride spectral power fluctuations than younger adults in sensorimotor and posterior parietal areas as terrain unevenness increases. This would relate to a higher demand for sensorimotor processing. We also performed a whole-brain exploratory analysis to reveal age differences in electrocortical activity. Findings from this study may uncover the underlying neural mechanisms that contribute to mobility limitations in older adults and inform targeted interventions to enhance mobility.

## Method

### Participants

This study is part of a larger multimodal brain imaging study (Mind in Motion), which investigates walking and mobility decline in older adults (NIH U01AG061389) (Clark et al., 2020). Here, we analyzed the cross-sectional EEG data from the larger study. We recruited 35 healthy younger (19 females, mean age = 24 ± 4 years, mean walking speed on uneven terrain treadmill = 0.7 +/-0.2 m/s) and 96 older (58 females, mean age = 75 ± 7 years, mean walking speed on uneven terrain treadmill = 0.35 ± 0.21 m/s) adults to assess their brain dynamics during walking on uneven terrain. Inclusion criteria consisted of younger adults being aged 20-40 years and community-dwelling older adults aged 65 years or older with a wide range of mobility. Exclusion criteria included the presence of cognitive impairments assessed by Montreal Cognitive Assessment [MoCA] score < 26, existing medical conditions that would significantly interfere with walking ability yet are not directly related to brain function, visual impairment that is not corrected, and implants with contraindications for Magnetic Resonance Imaging (MRI). Full inclusion and exclusion criteria have been described previously (Clark et al., 2020). This study was approved by the University of Florida Institutional Review Board (IRB 201802227). All participants provided written informed consent.

We included 31 younger adults and 71 older adults for data analysis (**Table 1**). A total of 3 younger adults were excluded from data analysis due to 1) gelling issues with braids (n = 1) and 2) technical issues (n = 2). 25 older adults were excluded for data analysis (**Supplementary** Figure 1) due to 1) not completing the MRI session (n = 8), 2) technical issue occurring during data collection (n = 1), 3) not completing High terrain conditions (n = 7), or 4) walking with a speed slower than 0.1 m/s because these participants tended to walk toward the front of the treadmill, pause their walking, and then let the treadmill transport them toward the back (n = 9).

**Table 1.** Participant characteristics, racial background/ethnicity for younger and older adults. Average (standard deviation) age, treadmill speed, short physical performance battery, 400-meter walking duration, 400-meter walking speed, and Montreal Cognitive Assessment for younger and older adults.

| Characteristics | Included Younger Adults (n = 31) | Included Older Adults (n = 71) | P value |
| --- | --- | --- | --- |
| Age (yrs), mean (SD) | 24(4) | 75(8) | n.a |
| Sex, M/F | 15/16 | 31/40 | n.a |
| Hand dominance, Right/Left/Both | 28/3 | 65/5/1 | n.a |
| Treadmill speed for all terrains (m/s), mean (SD) | 0.70(0.16) | 0.39(0.20) | P <.001 |
| Short Physical Performance Battery (score)<sup>a</sup>, mean (SD) | 12(0) | 9(1.7) | P <.001 |
| 400m walk (sec), mean (SD) | 326(40.5) | 389.5(63) | P <.001 |
| 400m speed (m/s), mean (SD) | 1.2(0.15) | 1.05(0.17) | P <.001 |
| Montreal Cognitive Assessment (score)<sup>a</sup>, mean (SD) | 28(1.4) | 27.5(1.6) | P = .06 |

| Racial background/Ethnicity | Included Younger Adults (n = 31) | Excluded Younger Adults (n = 4) | Included Older Adults (n = 71) | Excluded Older Adults (n = 25) |
| --- | --- | --- | --- | --- |
| White | 22 | 3 | 63 | 23 |
| Black | 1 | 1 | 3 | 2 |
| Asian | 6 | 0 | 2 | 0 |
| Other | 2 | 0 | 3 | 0 |
| Hispanic | 7 | 1 | 7 | 0 |

<sup>a</sup>These assessments were performed with 20 younger adults.

### Experimental protocol

Details of the study protocol have been previously reported (Clark et al., 2020; Liu et al., 2024) and we provide a summary of the procedures and setup below. We included one session of EEG and one session of MRI scans for this paper. The EEG and MRI sessions were conducted on separate days, with an average interval of 30 days between sessions (standard deviation = 91 days).

Prior to the EEG session, participants walked on an overground version of the Flat, Low, Medium, and High Terrain uneven conditions on a 3.5-meter mat three times. The overground speed for each terrain was computed as the average speed to walk through the middle 3-meter portion. During the EEG session, participants walked on the custom-designed treadmill belts at a subject-specific walking speed on four different levels of uneven terrain (Flat, Low, Med, and High) and at four different speeds (0.25 m/s, 0.5 m/s, 0.75 m/s, 1.0 m/s) on the flat surface. Subject-specific treadmill speed for all terrains was set to 75% of the slowest overground speed across the terrains unless participants requested a slower treadmill speed. Each condition consisted of two walking trials, each lasting three minutes. Participants also completed a three-minute seated resting trial (**Figure 1A**). During treadmill walking, participants wore a harness to prevent falling to the ground, but the harness did not provide any body weight support unless a fall occurred. No falls occurred during data collection. We pseudorandomized the conditions with 8 unique orders of uneven terrain conditions and speed conditions, respectively.

**Figure 1:**
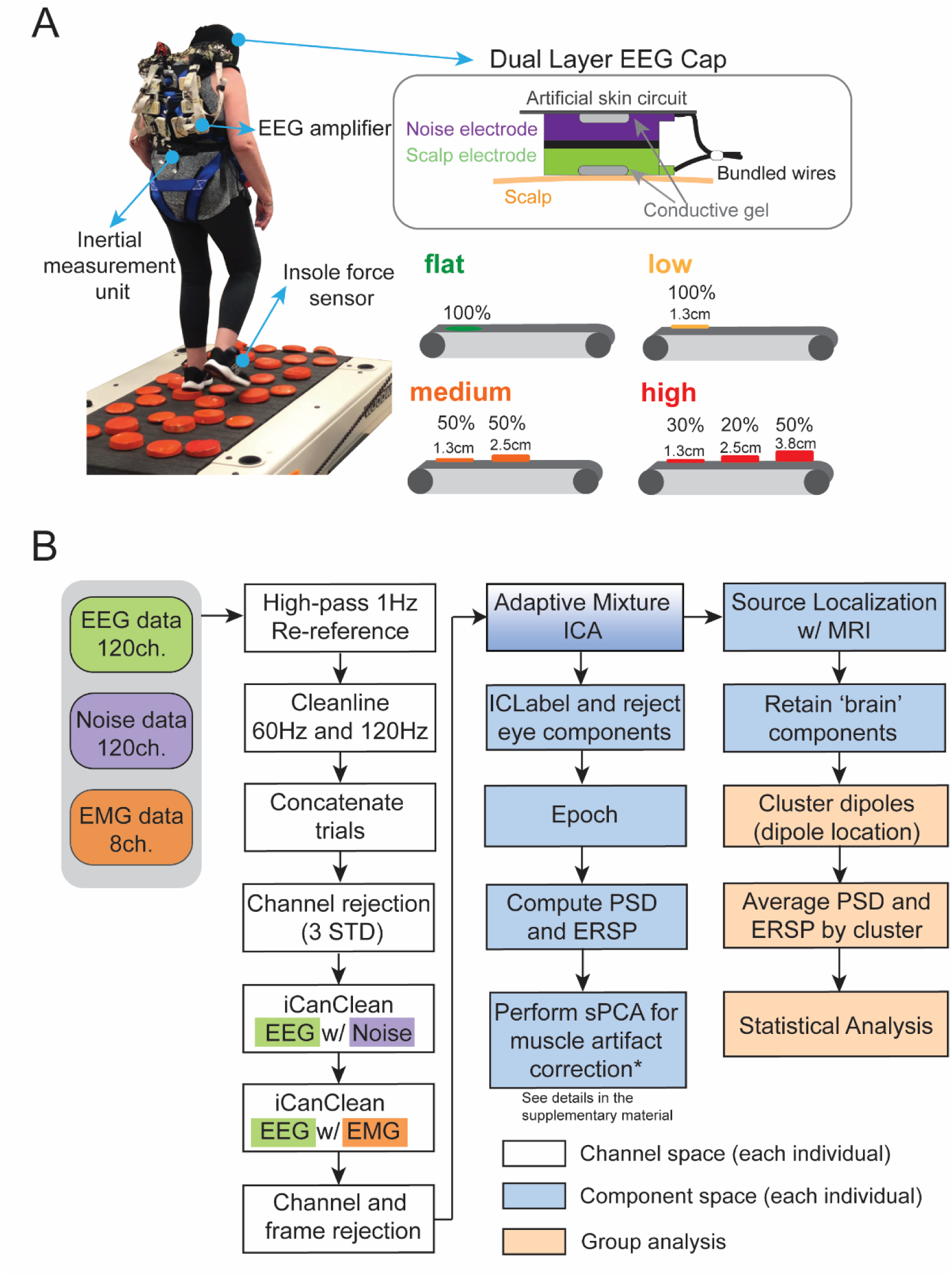
Experiment setup and processing pipeline. (A) Participants completed treadmill walking trials on four different levels of uneven terrain (Flat, Low, Medium, High) and other walking trials at different speeds performed on the Flat terrain. Participants wore a custom-made dual-layer EEG cap, an inertial measurement unit on their sacrum, and insole force sensors inserted in their shoes. Dual-electrode EEG setup has scalp electrodes and noise electrodes that were mechanically coupled. We used a conductive fabric as an artificial skin circuit to bridge the noise electrodes. (B) Data processing flowchart with steps for EEG pre-processing, source localization, spectral Principal Component Analysis, and clustering brain components.

### Data Acquisition

Participants wore a custom-made dual-layer EEG cap (ActiCAP snap sensors; Brain Products GmbH, Germany), including 120 scalp electrodes and 120 mechanically coupled noise electrodes, when walking on the treadmill. The scalp electrodes followed a 10-05 electrode system. We inverted and mechanically coupled noise electrodes to the scalp electrodes (**Figure 1A**) (Nordin et al., 2018; Studnicki et al., 2022). We used a conductive fabric as an artificial skin circuit and bridged the noise electrodes. Eight of the original 128 scalp electrodes (TP9, P9, PO9, O9, O10, PO10, P10, and TP10) were repurposed to measure muscle activity of the sternocleidomastoid and trapezius on the left and right sides. We kept all scalp electrode impedance values below 15 Kohm during the setup. Ground and reference electrodes were kept below 5 Kohm. We digitized the electrode locations using a structural scanner (ST01, Occipital Inc., San Francisco, CA). We used four LiveAmp 64 amplifiers and logged EEG data at 500 Hz.

The online reference and ground electrodes were at CPz and Fpz, respectively.

### Gait Event and Kinematics Analyses

We analyzed kinematic and kinetic data using MATLAB2020b (Mathworks, USA, RRID:SCR_001622) to compute behavioral variables. We defined foot strike and toe off based on ground reaction forces measured with the capacitive shoe insole sensors (loadsol 1-184 sensor, Novel Electronics Inc., St. Paul, MN, USA) with 20 N threshold (Downey et al., 2022). We computed the peak-to-peak excursion of the sacrum in the anteroposterior and mediolateral direction using the IMU. We have previously reported the details of the algorithm and calculations (Downey et al., 2022). We removed outliers that were +/-2.5 standard deviations away from the mean within each trial (Downey et al., 2022). We calculated the variability of each of these measures as the coefficient of variation (standard deviation over mean).

### EEG Data Analyses

We processed all EEG data using custom MATLAB scripts, EEGLAB (v 2021.0, RRID:SCR_007292) (Delorme & Makeig, 2004), and the BeMoBIL pipeline (v2.0.0) (Klug et al., 2022) (**Figure 1B**). We applied a 1 Hz high-pass filter (−6 dB at 0.5 Hz) with *eegfiltnew* on all scalp, noise, and muscle channels to remove drift for each trial and then applied a 20 Hz high-pass filter with *eegfiltnew* on muscle channels. We used the CleanLine plugin in EEGLAB to remove line noise at 60 Hz and 120 Hz. We rejected bad channels that were 3 standard deviations away from the mean of EEG and noise channels, respectively. We performed average reference for scalp, noise, and muscle channels respectively. We then used *iCanClean* (Downey & Ferris, 2023; Gonsisko et al., 2023) to remove artifacts that were highly correlated with noise reference electrodes (R^2^ = 0.65 with a four-second moving window) and muscle reference electrodes (R^2^ = 0.4 with a four-second moving window). We used *clean_artifacts* in EEGLAB to remove bad channels and noisy time frames using default parameters except for the following: chan_crit1 = 0.7, win_crit1 = 0.4, winTol = [-Inf, 10]. These parameters were selected in a preliminary analysis of a subset of the data, which aimed to minimize the number of channels and time frames rejected while maximizing a good number of brain components by ICLabel (Liu et al., 2023; Pion-Tonachini et al., 2019). We retained 110 +/-6 channels after pre-processing. Scalp EEG data were re-referenced. We performed adaptive mixture independent component analysis (AMICA) on the preprocessed data to decompose the preprocessed EEG data into statistically independent components (Palmer et al., 2011). For this analysis, we used all EEG data (approximately 50 minutes), which included both terrain and speed walking trials, as well as the resting trial, for AMICA; however, we only used the terrain trials in subsequent data analysis. We later used the independent components to perform source localization.

### Epoch and compute power spectral density and event-related spectral perturbations

For the walking trials, we segmented data into epochs of 5.25 seconds (from 1 second before to 4.25 seconds after the right foot strike). The epoch length was chosen to accommodate participants with long step durations during the slowest walking condition (0.25 m/s). We rejected epochs that were three standard deviations from the mean gait event time. For the resting seated trial, we also segmented data into epochs of 5.25 seconds.

We computed the log power spectral density (PSD) using *spectopo* from EEGLAB with default parameters for each independent component and normalized by subtracting each individual’s mean log spectral power density from all conditions. We then computed the time-frequency decomposition of the resting trial with *newtimef* (Morlet wavelets cycles: [3 0.8]) and averaged across all epochs and time. We also computed the time-frequency decomposition for all walking trials and then time-warped the gait cycles from right foot-strike, left foot-off, left foot-strike, right foot-off, and the subsequent right foot-strike. We then averaged across all epochs. We calculated relative changes in power between walking and resting conditions of every time point in the gait at every frequency for each independent component.

### Muscle artifact correction with sPCA

We used an adapted spectral principal component analysis (sPCA; N. S. J. Jacobsen et al., 2022; Seeber et al., 2015) to obtain muscle artifacts for correction in event-related spectral perturbations (ERSP) and power spectral density (PSD) for each participant. We performed sPCA for additional muscle artifact reduction in addition to iCanClean because there were residual muscle artifact contaminations found in the frequency domain, especially for older adults. The steps were previously described in Salminen et al.(2025) in detail, and thus we provided a brief summary here and in the **Supplementary** Figures 2 **-3**. We followed the scripts provided by Jacobsen et al. to implement the sPCA (N. S. J. Jacobsen et al., 2022). For each participant, we first removed the eye components identified by ICLabel (Pion-Tonachini et al., 2019) (version: lite). We computed the relative changes in PSDs and ERSPs during all walking trials from the resting trial for each component. sPCA was performed on those PSDs and ERSPs averaged across conditions. We obtained eigenvectors and a weighting matrix that transformed data from component x frequency space to the principal component space. The first principal component (PSC1) with the largest eigenvalue was identified as muscle artifacts and thus removed from further analysis. Similar to Seeber et al. (2015), the spectra profile of the PSC1 was lower at frequency 2 – 20Hz but became much higher at higher frequencies, suggesting that PSC1 reflected electromuscular activity (**Supplementary** Figure 4). We then back-projected the remaining principal components using the weighting matrix for each condition (Seeber et al., 2015). We used the same weighting matrix for all conditions for each participant. After obtaining the sPCA-corrected PSD, we added the PSD from the resting condition to the sPCA-corrected PSD (**Supplementary** Figure 2; Salminen et al., 2025) to allow the FOOOF (Fitting oscillations & one over f) toolbox to separate the aperiodic and periodic components (Donoghue et al., 2020).

**Participant-specific volume conduction head model and source localization** We created the participant-specific volume conduction head model using each participant’s T1-weighted MRI with Fieldtrip (v. 20210910, RRID:SCR_004849). After we resliced the image, we performed tissue segmentation using headreco from SimNIBS toolbox (v 3.2). The MRIs were segmented into six tissue layers (scalp, skull, air, cerebrospinal fluid, gray matter, and white matter). Then we generated finite element hexahedral meshes following the steps previously reported in Liu et al. 2023.

We digitized the fiducial locations (left/right preauricular, nasion) on the MRI and then co-registered the digitized electrode locations to the individual-specific head models by aligning the fiducial locations. We then computed the leadfield matrix using SIMBIO toolbox with a 5 mm apart distributed source position in gray matter.

We performed EEG source localization with an equivalent dipole fitting approach using *ft_dipolefitting* function in the Fieldtrip toolbox. We then warped the dipole locations to the Montreal Neurological Institute (MNI) template for both younger and older adults using ANTs normalization (Advanced Normalization Tools, https://github.com/ANTsX/ANTs, RRID:SCR_004757, Avants et al., 2011). The dipole locations found in the subject-specific head model were warped to the MNI template using *antsApplyTransformsToPoints.* We retained brain components using the following criteria: 1) ICLabel (Pion-Tonachini et al., 2019) (version: lite) classified the brain probability of greater or equal to 50%, 2) negative slope of the power density spectrum for 2 - 40 Hz to remove muscle components, 3) residual variance of dipole fitting <15%, and 4) dipoles located inside the brain. Ten older adult participants with fewer than 5 brain components were removed from further analysis. The number of remaining brain components was greater in younger adults than older adults (15 +/-5 vs. 12 +/-5, t(88) = 2.5, p = 0.01).

### K-means clustering of brain components

We clustered the brain components into 11 clusters (k = 11) by dipole locations using robust k-means (maxiter = 10,000 and replicate = 1000) in EEGLAB. We used silhouette, Calinski-Harabasz, and Davies-Bouldin methods to evaluate the clustering outcome between the range of 9 – 14 clusters, as 14 is the average number of brain components across participants. We did not find agreement when evaluating the clustering outcome between k = [9, 14] using the three evaluation methods. We chose k = 11 so that the cluster locations would be the most comparable to our previous paper with young adults (Liu et al., 2024). Clusters with at least half of the younger adults (n >= 16) and half of the older adults (n >= 30) were retained. Components that were three standard deviations away from any of the cluster centroid were identified as outliers. Each cluster had at most one brain component from each participant. We selected the component with the maximum likelihood to be a brain component by ICLabel to prevent inflating the sample size if multiple components per subject existed in a cluster.

### Averaging power spectral density and normalizing event-related spectral perturbations for each cluster

For each cluster, we averaged the power spectral density and event-related spectral perturbations after sPCA muscle artifact correction. We used the FOOOF toolbox to separate the aperiodic and periodic components from the power spectral densities (Donoghue et al., 2020). We set the FOOOF parameters as follows: range of power spectra [3 40 Hz], peak width limits: [1 8], minimum peak height: 0.05, maximum number of peaks: 2. We computed the average power for each band using the flattened power spectral density by subtracting the aperiodic component from each of the original power spectral density.

We computed ERSP with two different normalization methods. First, we normalized ERSPs to the average spectral power across gait cycles within each condition to investigate the power fluctuation within gait cycles or intra-stride power fluctuations. We also obtained the peak-to-peak ERSP for each band (theta, alpha, beta) for each terrain condition by taking the average of the power across all frequencies within each band throughout the gait cycle and then computing the peak-to-peak range for each terrain condition. On the other hand, we normalized ERSPs to average spectral power across gait cycles for all conditions (common baseline removal) to allow for comparison of ERSPs across terrain conditions. We averaged the ERSPs for each terrain condition and then averaged across all participants for each cluster.

### Statistical Analysis

All statistical analyses were performed in MATLAB 2020b (Mathworks) or Rstudio (v4.4) for linear mixed effect models. All significance levels (α) were set at 0.05. False discovery rate (FDR) controlling procedures were implemented to control for all multiple comparisons (Benjamini & Hochberg, 1995). We first assessed the participant characteristic differences between younger and older adults using Welch t-tests for normally distributed continuous data, Wilcoxon Rank Sum Test for non-normally distributed continuous data, and chi-square (χ^2^) test for categorical data.

We used R (package: *lme4, lmerTest, emmeans*) for all linear mixed-effect analyses for behavior outcomes, as in our previous paper (Downey et al., 2022). We first assessed whether any of the behavioral measures were affected by terrain and age. The dependent variables included the step duration coefficient of variation and the sacral excursion coefficient of variation in mediolateral and anteroposterior directions. The independent variables included Terrain (Flat, Low, Medium, High), Age group (Younger adults and Older adults), and their interaction. Walking speed was included as a covariate for all mixed-effect models to control for the potential confounding effect. We included a random intercept to account for unmodeled sources of between-subject variability. Post hoc analysis was performed if we found a significant main or interaction effect using ANOVA (package: *car*). Effect size was reported using partial eta-squared (η²ᵖ) with 0.01 for small effect size, 0.06 for medium effect size, and 0.14 for large effect size. If the interaction was insignificant, we refit a linear mixed-effect model with no interaction. Post hoc analyses were performed by setting up contrast matrices and pairwise comparisons using *emmeans*. False discovery rate (FDR) controlling procedures were implemented to control for multiple comparisons (Benjamini & Hochberg, 1995).

We used MATLAB for EEG power spectral and gait-related spectral power fluctuation statistical tests. We assessed if the power spectral density for each cluster differed across terrain and age groups. We performed non-parametric permutation statistics for flattened power spectral density using Fieldtrip in EEGLAB (α = 0.05, 2000 iterations) and corrected for multiple comparisons across frequency using a false discovery rate at 0.05. The non-parametric permutation statistics could not assess the interaction effect between terrain and age group due to the inherent limitation of this test. We evaluated the within-stride gait-related spectral power for each terrain condition and for each age group. These ERSPs were normalized to the average spectral power across all gait cycles within conditions. We performed bootstrapping (α = 0.05, 4000 iterations) for each terrain condition in each brain cluster and corrected for multiple comparisons across time-frequency using a false discovery rate at 0.05.

We used MATLAB to evaluate gait-related spectral power during different terrain conditions and examined differences between younger and older adults. We first quantified gait-related spectral power during uneven terrain walking with respect to the Flat condition for younger and older adults, respectively (ΔERSP = ERSP_uneven_ – ERSP_Flat_). We then quantified the difference in gait-related spectral power between younger and older adults by subtracting the younger group’s gait-related spectral plot from the older group’s gait-related spectral plot (ΔERSP_young_ – ΔERSP_old_). We conducted statistical analyses using cluster-based permutation tests (α = 0.05, 10,000 permutations) calculated with FieldTrip functions within EEGLAB (Delorme & Makeig, 2004; Maris & Oostenveld, 2007), which uses the Monte Carlo method to estimate permutation p-values. Cluster-based permutation testing is a robust nonparametric statistical approach that corrects for multiple comparisons across time-frequency and reduces the potential for false negatives in high-dimensional EEG data (Maris & Oostenveld, 2007).

We also used R for all linear mixed-effect analyses to assess whether the average power spectral density after FOOOF for each band (theta, alpha, and beta) and peak-to-peak within-stride spectral power fluctuations were affected by terrain and age group for each brain cluster and each band. Outliers more than 8 standard deviations from the mean were excluded from the analysis. To account for the effect of each individual’s preferred treadmill speed used on uneven terrain on our outcome measures, we first tested whether average power or peak-to-peak power fluctuations were associated with treadmill speed. Since we did not find a significant association between most EEG outcome measures and treadmill speed, we only included treadmill speed as a covariate in all subsequent models. The independent variables included Terrain (Flat, Low, Med, High), Age group (Younger adults and Older adults), and their interaction.

We included a random intercept for each model. The Flat condition was set as the reference for younger adults. We performed post hoc analysis if we found a significant main or interaction effect using ANOVA (package: *car*). Effect size was reported using partial eta-squared (η²ᵖ). If the interaction was insignificant for a given band at the given brain area, we refit a linear mixed-effect model with no interaction. Post hoc analyses were performed by setting up contrast matrices and pairwise comparisons using *emmeans*. False discovery rate (FDR) controlling procedures are implemented to control for between-condition and between-group multiple comparisons (Benjamini & Hochberg, 1995).

## Results

### Participant characteristics

Older adults walked significantly slower than younger adults on the uneven terrain treadmill (**Figure 2A**, **Table 1**). The slower treadmill speed for older adults was due to their slower overground walking speed on the uneven terrain mat. Older adults also walked slower during the 400-meter test than younger adults (**Table 1**) and scored lower on the short physical performance battery test compared to younger adults (**Table 1**). There were no differences in MoCA between younger and older adults. Excluded older adult participants were, on average, three years older than those who were included, likely because many of the excluded individuals who had walking speeds slower than 0.1 m/s were older. There was no significant difference in sex distribution between included and excluded participants.

**Figure 2:**
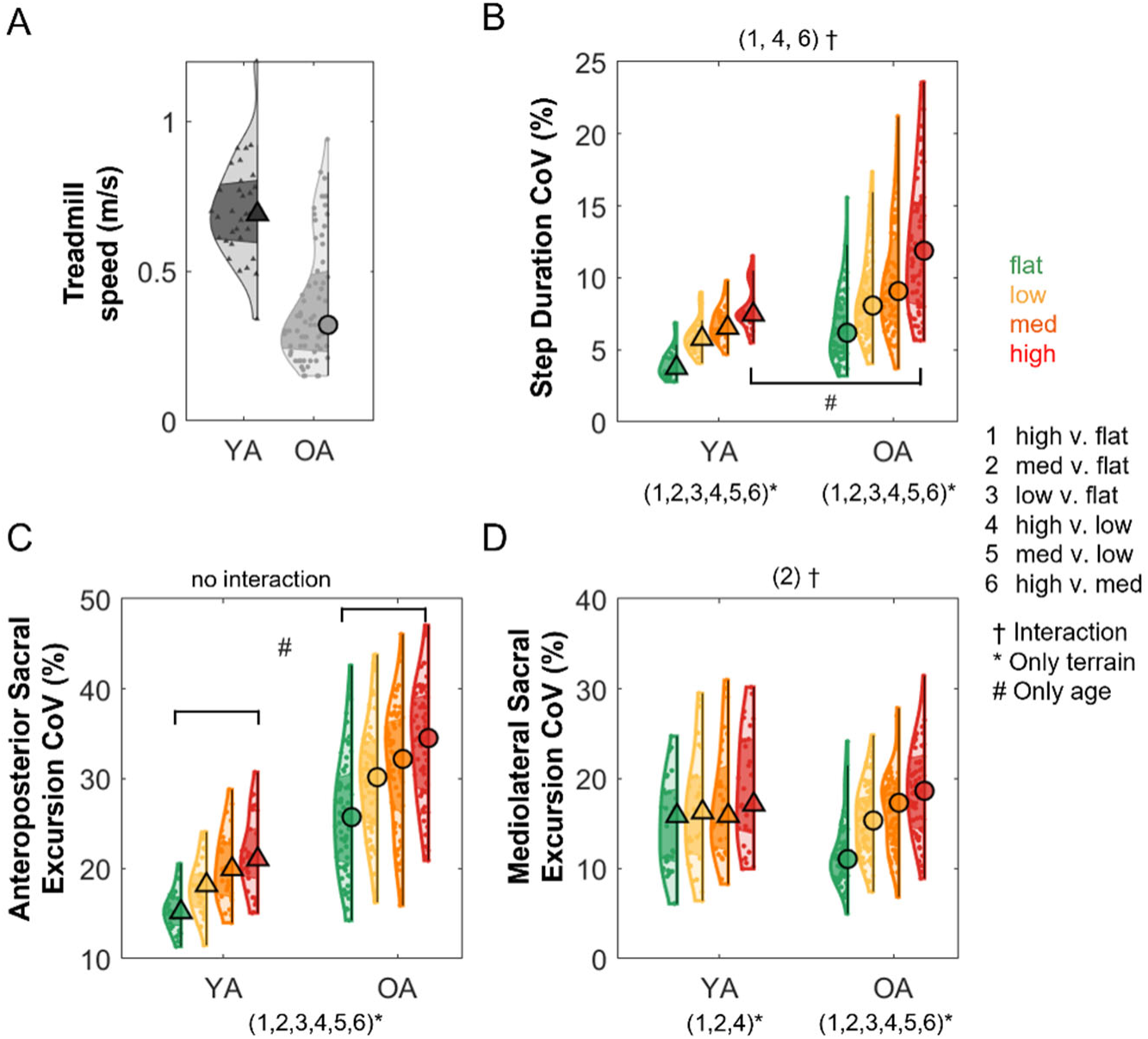
Violin plots show the behavioral measures during walking on the uneven terrain treadmill. (A) Uneven terrain treadmill walking speed for younger adults (YA) and older adults (OA). (B) Step duration coefficient of variation (CoV) at different terrains. Sacral excursion coefficient of variation (CoV) in the anteroposterior (C) and mediolateral (D) direction. The shaded regions represent data distribution across participants by estimating the probability density function. The triangle and circle markers represent the median of the data. The darker shaded region represents the 25th to 75th percentiles of the data. †: significant interactions, *: significant comparison between terrain conditions within each group, # significant age group effect for a specific terrain condition.

### Behavior measures

We only provide a summary of behavioral measures here. All these behavioral results have been previously reported in our previous paper with a smaller subset of the participants (Downey et al., 2022). The current study extends those findings by including the full participant sample recruited for the Mind in Motion study. We reported the variability measures here as they have high validity for assessing gait stability (Bruijn et al., 2013). Variability was computed as the coefficient of variation (standard deviation over mean), and walking speed was added as a covariate to control for the confounding effect for all statistical analyses. Walking on an uneven terrain treadmill increased step duration coefficient of variation and sacral excursion coefficient of variation in both anteroposterior and mediolateral directions after accounting for walking speed (**Figure 2, Supplementary Table 1**). Both older adults and younger adults increased their step duration coefficient of variation with terrain unevenness (all p_FDR_ < 0.05), and older adults showed a greater increase in step duration between the High terrain vs. Flat terrain (p_FDR_ = 0.003), High terrain vs. Low terrain (p_FDR_ = 0.006), and High terrain vs. Med terrain (p_FDR_ = 0.027) conditions. Both groups also increased the sacral excursion coefficient of variation in the anteroposterior direction (all p_FDR_ < 0.001), with older adults demonstrating greater coefficient of variation in all conditions (p < 0.001). Older adults increased the sacral excursion coefficient of variation in the mediolateral direction with terrain unevenness (all p_FDR_ < 0.01), while younger adults only increased the coefficient of variation between the Med terrain vs. Flat terrain (p_FDR_ < 0.002), High terrain vs. Flat terrain (p_FDR_ < 0.001), and High terrain vs. Low terrain (p_FDR_ = 0.006). Older adults also showed a greater increase in the mediolateral sacral excursion coefficient of variation in the Med terrain vs. Flat terrain than younger adults (p_FDR_ = 0.004). Full statistical results, including multiple comparisons, are provided in **Supplementary Table 1-4**. In summary, older adults demonstrated a greater increase in step duration variability and mediolateral sacral excursion variability with terrain unevenness than younger adults.

### EEG source analysis

We identified nine brain source clusters (**Figure 3**; **Table 2**). The dipole clusters were located at the left sensorimotor, right sensorimotor, left posterior parietal, right posterior parietal, left pre-supplementary motor, right premotor, precuneus, mid cingulate, and temporal area. The temporal area was excluded from further analysis, as we did not have a hypothesis related to this area.

**Figure 3:**
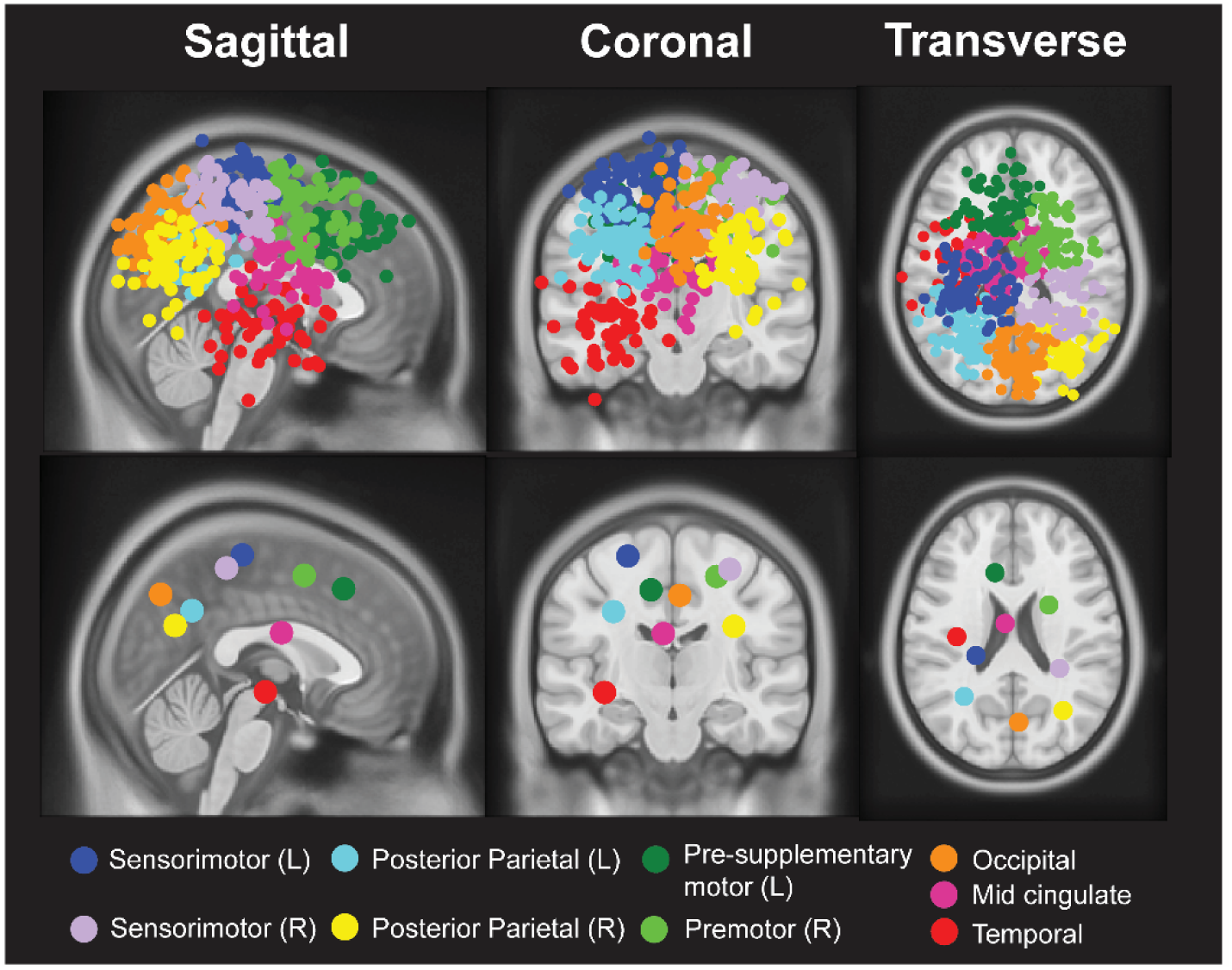
Dipole location for all participants (top row) and centroid of each cluster (bottom row) in sagittal, coronal, and axial planes. We identified clusters located at: right sensorimotor area, left sensorimotor area, left posterior parietal, right posterior parietal, occipital area, mid cingulate, left pre-supplementary motor, right premotor, and left temporal area. Brain sources in this figure represent spatial localization defined by ICA methods rather than the brain sources that showed differences across uneven terrain conditions.

**Table 2:** Montreal Neurological Institute (MNI) coordinates, and anatomical atlas labels for regions of interest (ROIs).

| Cluster centroid location | Color | No. participants | MNI coordinates | Anatomical Label of the centroid |
| --- | --- | --- | --- | --- |
| Sensorimotor (L) | Blue | 27 younger/49 older | -23 -29 64 | Postcentral L<sup>a</sup> |
| Sensorimotor (R) | Purple | 23 younger/41 older | 30 -37 57 | Postcentral R<sup>a</sup> |
| Posterior Parietal (L) | Aqua | 27 younger/43 older | -31 -55 35 | Angular L<sup>a</sup> |
| Posterior Parietal (R) | Yellow | 25 younger/33 older | 32 -64 28 | No label<sup>a</sup> |
| Occipital | Orange | 26 younger/50 older | 4 -71 44 | Precuneus<sup>a</sup> |
| Mid cingulate | Pink | 21 younger/39 older | -5 -8 24 | No label<sup>a</sup> |
| Pre-supplementary Motor (L) | Dark Green | 24 younger/35 older | -11 24 47 | Frontal_Sup_L<sup>a</sup> |
| Premotor (R) | Lime | 22 younger/38 older | 23 3 54 | Frontal_Sup_R<sup>a</sup><br>Premotor<sup>b</sup> |
| Temporal (L) | Red | 19 younger/32 older | -36 -17 -7 | No Label<sup>a</sup> |

<sup>a</sup> Anatomical location of the cluster centroid was labeled based on (Tzourio-Mazoyer et al., 2002).
<sup>b</sup> Anatomical location of the cluster centroid was labeled based on (Mayka et al., 2006).

### EEG power spectral density

We found significant effects of terrain unevenness (p < 0.05) on EEG power spectral density at all brain clusters using non-parametric permutation statistics (**Figure 4, Supplementary** Figure 5). Theta band power was higher with increased terrain unevenness in the sensorimotor, posterior parietal, and occipital areas. Alpha band power was lower with increased terrain unevenness in bilateral sensorimotor, bilateral posterior parietal, mid cingulate, occipital area, and right premotor area. We also observed lower beta band power with increased terrain unevenness in all brain clusters.

**Figure 4:**
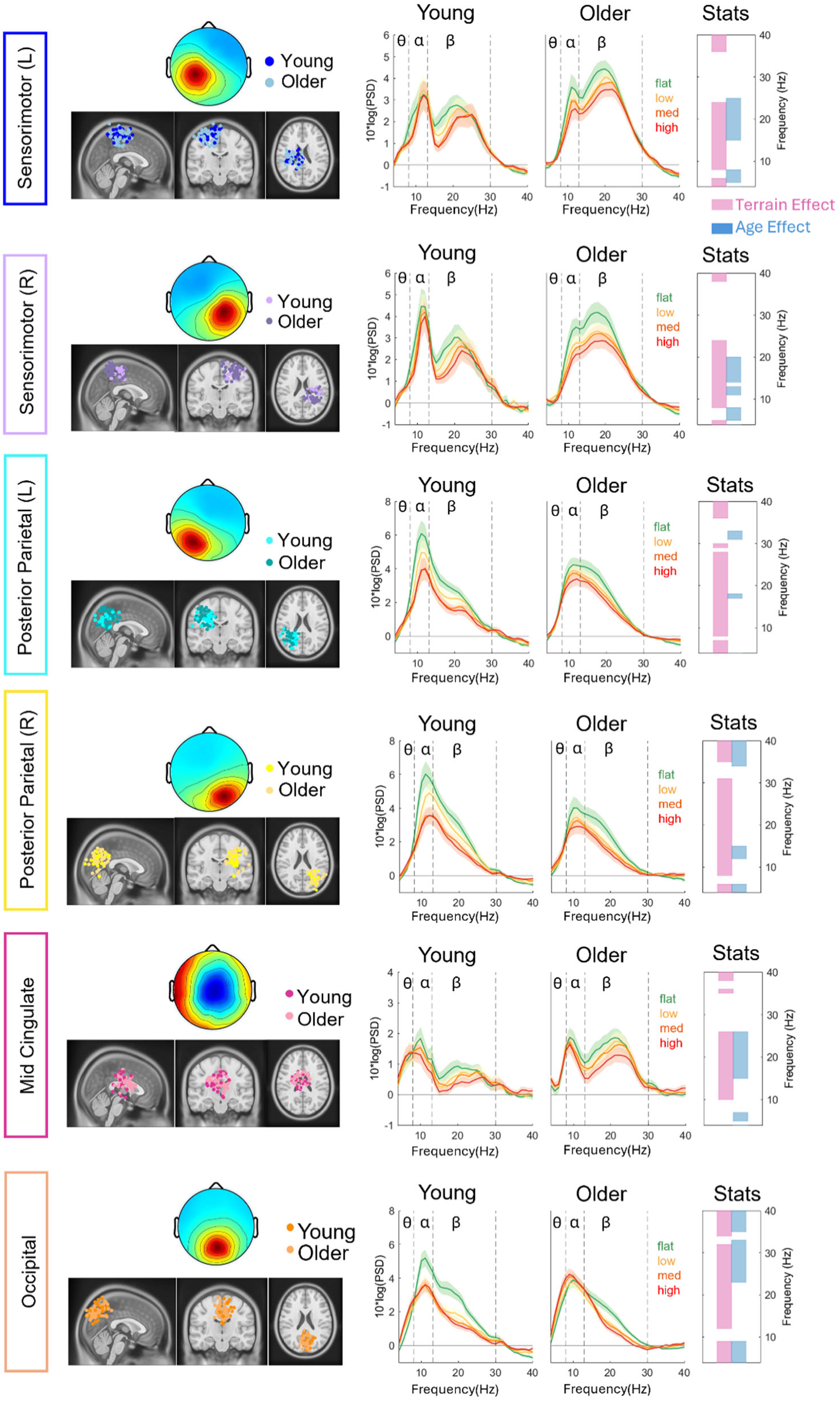
Dipole location plotted on the Montreal Neurological Institute template, scalp topography, and average flattened power spectrum densities (PSDs) changed with terrain unevenness for younger and older adults for bilateral sensorimotor clusters, bilateral posterior parietal clusters, mid cingulate, and occipital cluster. Shaded colored areas indicate standard error of PSDs across components in the cluster. Vertical black dashed lines indicate main frequency bands of interest—theta (4 - 8 Hz), alpha (8 - 13 Hz), and beta (13 - 30 Hz). The very right panel indicates the significant terrain and age effect on PSD at each frequency (pink: terrain, blue: age). In this non-parametric statistical analysis, only main effects were tested due to the limitation of this test.

We also found significant effects of age (p < 0.05) in the bilateral sensorimotor, right posterior parietal, occipital, and mid cingulate clusters (**Figure 4, Supplementary** Figure 5). Pooled theta band power was generally greater in younger adults than older adults in bilateral sensorimotor, right posterior parietal, and mid cingulate cluster, but it was the opposite in the occipital cluster. Alpha band power was greater in younger adults than older adults in the right sensorimotor cluster and right posterior parietal cluster. Lastly, beta band power was greater in older adults than younger adults in the sensorimotor and mid-cingulate cluster, but it was the opposite in the occipital cluster.

### Age and terrain unevenness on average EEG power

We tested whether average theta power, alpha power, and beta power were affected by age, terrain, and interaction for each cluster after controlling for treadmill walking speed (**Figure 5**). Full statistical analysis results are reported in **Supplementary Table 5-12**.

**Figure 5:**
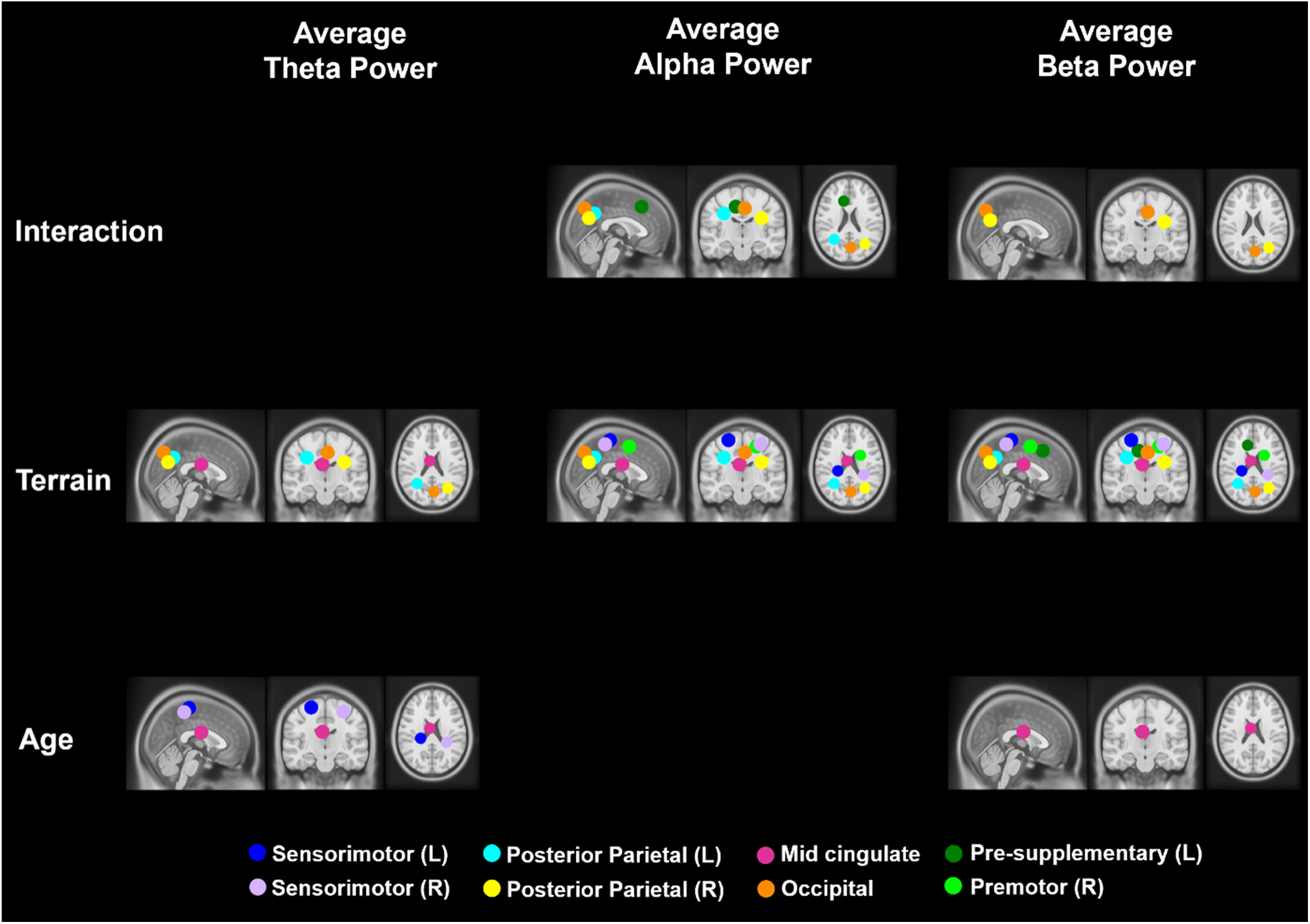
Significant effects of age, terrain, and interaction on theta, alpha, and beta average power (p < 0.05). The analyses were controlled with the effect of treadmill walking speed. Only centroids of each cluster where significant results were found are plotted.

**Interaction effect**: The effect of terrain on EEG alpha power depends on the age group in the parietal-occipital region (**Figure 6**). When walking on uneven terrain, younger adults exhibited a significant reduction in alpha power compared to walking on a flat surface, whereas older adults showed little to no reduction. An interaction effect of age and terrain on **average alpha power** was found in the bilateral posterior parietal (Left: F(3,204) = 9.26, p < 0.001, η²ᵖ = 0.12, Right: F(3, 168) = 3.63, p = 0.014, η²ᵖ = 0.061) and occipital clusters (F(3, 222) = 5.86, p < 0.001, η²ᵖ = 0.073; **Figure 6A**). A main effect of terrain was also found in these clusters (all p < 0.001). At the <u>left posterior</u> <u>parietal</u> cluster, older adults had less alpha power reduction compared to younger adults in the High terrain versus Flat (p_FDR_ < 0.001), Med terrain versus Flat (p_FDR_ < 0.001), and Low terrain versus Flat (p_FDR_ = 0.017). Additionally, younger adults exhibited alpha reduction at greater terrain unevenness in all pairwise comparisons (all p_FDR_ < 0.01) except the High terrain versus Med, while older adults only exhibited alpha reduction at High terrain versus Flat (p_FDR_ = 0.001). At the <u>right posterior parietal</u> cluster, older adults had less alpha power reduction in the High terrain versus Flat (p_FDR_ = 0.036) and Med terrain versus Flat (p_FDR_ = 0.016) compared to younger adults. Younger adults exhibited significant alpha reduction at High terrain versus Flat, Med terrain versus Flat, and Low terrain versus Flat (all p_FDR_ < 0.001). Older adults only exhibited alpha reduction at High terrain versus Flat (p_FDR_ = 0.022). At the <u>occipital cluster</u>, older adults had less difference in alpha power reduction in the High terrain versus Flat (p_FDR_ = 0.001), Med terrain versus Flat (p_FDR_ = 0.002), and Low terrain versus Flat (p_FDR_ = 0.027) compared to younger adults. Younger adults exhibited significant alpha reduction at High terrain versus Flat, Med terrain versus Flat, and Low terrain versus Flat (all p_FDR_ < 0.01), but no alpha reduction was found in older adults.

**Figure 6:**
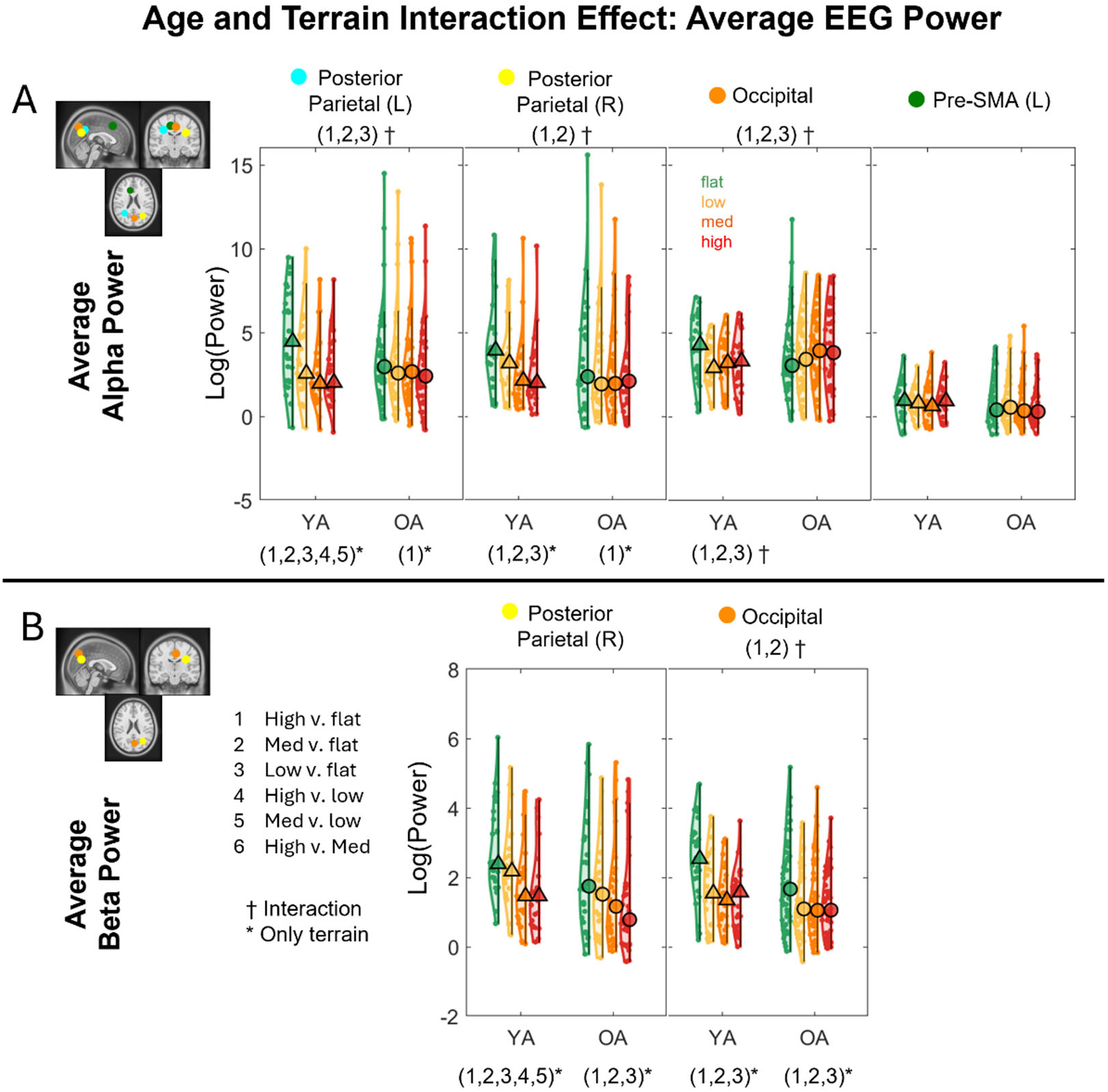
Violin plots of the average alpha power (A) and beta power (B) for each terrain and age group for the corresponding brain clusters with a significant age and terrain interaction effect. YA: younger adults, OA: older adults. #Indicates significant group differences between two terrain conditions, *Indicates significant differences between terrain conditions. †Indicates significant interaction effects between terrain and age group. 1: High vs. Flat, 2: Med vs. Flat, 3: Low vs. Flat, 4: High vs. Low, 5: Med vs. Low, 6: High vs. Med. Circle and triangle markers indicate median across participants.

While the left pre-supplementary cluster also had significant interaction (F(3, 171) = 2.83, p = 0.04), none of the pairwise comparisons was significant after false discovery rate adjustment.

The effect of terrain on EEG beta power depends on the age group in the parietal-occipital region (**Figure 6B**). The interaction effect of age and terrain on **average beta power** was found in the right posterior parietal (F(3, 168) = 3.38, p = 0.02, η²ᵖ = 0.057) and occipital clusters (F(3, 222) = 6.36, p < 0.001, η²ᵖ = 0.079). The main effect of terrain was also found in these clusters (both p < 0.001). At the <u>right posterior parietal</u> <u>cluster</u>, younger adults exhibited beta power reduction at greater terrain unevenness in all pairwise comparisons (all p_FDR_ < 0.001) except for High terrain versus Med terrain, while older adults exhibited beta power reduction only at High terrain versus Flat, Med terrain versus Flat, and Low terrain versus Flat (p_FDR_ < 0.001). At the <u>occipital cluster</u>, older adults had less difference in beta band power reduction at High terrain versus Flat (p_FDR_ = 0.002) and Med terrain versus Flat (p_FDR_ < 0.001) compared to younger adults. Additionally, both younger and older adults exhibited beta power reduction at High versus Flat, Med versus Flat, and Low versus Flat (p_FDR_ < 0.001).

**Age effect**: Main effects of age on average theta band power were found in the sensorimotor clusters (Left: F(1,73) = 5.25, p = 0.025, η²ᵖ = 0.067; Right: F(1, 61) = 5.85, p = 0.019, η²ᵖ = 0.087), and mid cingulate cluster (F(1,57) = 5.38, p = 0.024, η²ᵖ = 0.086) with older adults having lower theta band power than younger adults after controlling for walking speed **(Supplementary** Figure 6A). Main effect of age on average beta band power was found in the mid-cingulate cluster (F(1,57) = 4.71, p = 0.034, η²ᵖ = 0.076) with older adults having greater beta power than younger adults **(Supplementary** Figure 6B).

**Terrain effect only**: In the absence of a significant interaction effect for a given brain cluster, we reran our linear mixed effect models to include only terrain and age group as main effects. A main terrain effect on the **average theta power** was found in the bilateral posterior parietal (Left: F(3, 207) = 5.86, p = 0.001, η²ᵖ = 0.078; Right: F(3, 171) = 3.97, p = 0.009, η²ᵖ = 0.065), mid cingulate (F(3, 177) = 3.59, p = 0.015, η²ᵖ = 0.057), and occipital cluster (F(3,225) = 29.46, p < 0.001, η²ᵖ = 0.282; **Figure 7A**). Greater theta power was associated with greater terrain unevenness. Both younger and older adult groups had greater theta power in High terrain vs. Flat (p_FDR_ = 0.008), High terrain vs.

**Figure 7:**
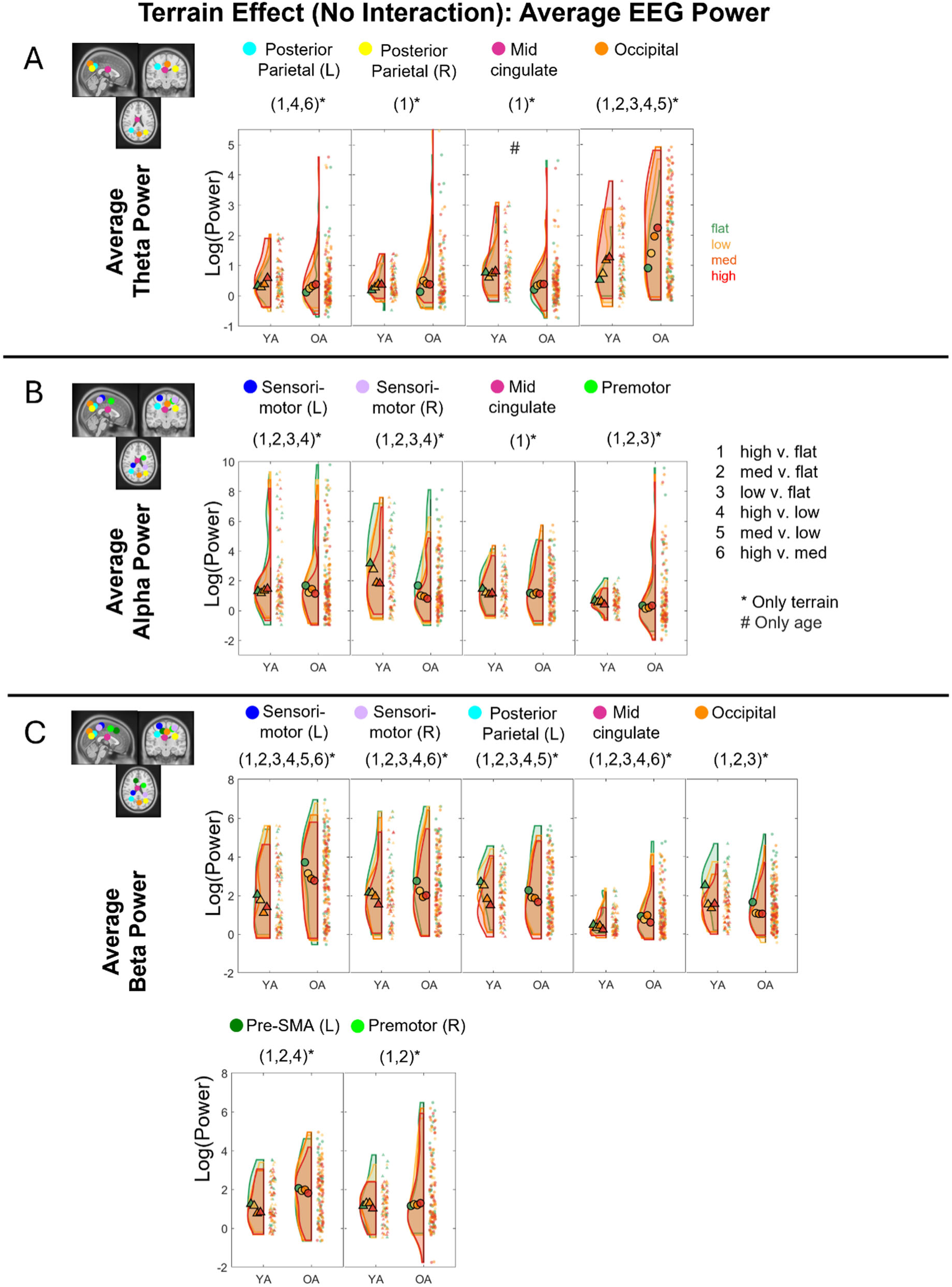
Raincloud plots of the average theta power (A), alpha power (B), and beta (C) power for each terrain and age group for the corresponding brain clusters without an interaction effect. YA: younger adults, OA: older adults. *indicates significant differences (p < 0.05) between terrain conditions with 1: High vs. Flat, 2: Med vs. Flat, 3: Low vs. Flat, 4: High vs. Low, 5: Med vs. Low, 6: High vs. Med. Circle and triangle markers indicate median across participants. # significant comparison between age groups for a specific terrain condition.

Low (p_FDR_ =0.001), and High terrain vs. Med (p_FDR_ = 0.044) terrain at the left posterior parietal area. Both groups showed greater theta power in High terrain vs. Flat (p_FDR_ = 0.007) terrain at the right posterior parietal, High terrain vs. Flat (p_FDR_ = 0.021) at the mid cingulate cluster. Both younger and older adult groups also showed significantly greater theta power at the occipital cluster during uneven terrain walking for all levels of comparisons (p_FDR_ < 0.01) except between High vs. Med terrain.

A terrain main effect on the **average alpha power** was found in the bilateral sensorimotor (Left: F(3, 225) = 18.31, p < 0.001, η²ᵖ = 0.20, Right: F(3, 189) = 18.85, p < 0.001, η²ᵖ = 0.23), mid cingulate (F(3, 177) = 4.16, p = 0.007, η²ᵖ = 0.066), and right premotor (F(3, 177) = 5.7, p = 0.001, η²ᵖ = 0.088) clusters (**Figure 7B**). Lower alpha power was associated with greater terrain unevenness. Both younger and older adults had significantly lower alpha power in High terrain vs. Flat (p_FDR_ < 0.001), Med terrain vs. Flat (p_FDR_ < 0.001), Low terrain vs. Flat (p_FDR_ < 0.01), and High terrain vs. Low (p_FDR_ < 0.01) terrain in bilateral sensorimotor areas. Both age groups also had lower alpha power in High terrain vs. Flat (p_FDR_ = 0.005) in mid-cingulate area and lower alpha power in High terrain vs. Flat (p_FDR_ = 0.001), Med terrain vs. Flat (p_FDR_ = 0.022), and Low terrain vs. Flat (p_FDR_ = 0.034) in the right premotor area.

Main terrain effect on **average beta power** was found in the bilateral sensorimotor (Left: F(3, 225) = 27.42, p < 0.001, η²ᵖ = 0.268; Right: F(3, 189) = 19.14, p < 0.001, η²ᵖ = 0.233), bilateral posterior parietal (Left: F(3, 207) = 37.37, p < 0.001, η²ᵖ = 0.351; Right: F(3, 168) = 43.32, p < 0.001, η²ᵖ = 0.436), mid-cingulate (F(3, 177) = 20.81, p < 0.001, η²ᵖ = 0.261), left pre-supplementary (F(3, 174) = 8.64, p < 0.001, η²ᵖ = 0.13), and right premotor (F(3, 177) = 7.86, p < 0.001, η²ᵖ = 0.118) clusters (**Figure 7C**). Lower beta power was associated with greater terrain unevenness. At <u>bilateral sensorimotor</u> <u>clusters</u>, both younger and older adults had significantly lower beta power in High terrain vs. Flat (both p_FDR_ < 0.001), Med terrain vs. Flat (both p_FDR_ < 0.001), Low terrain vs. Flat (both p_FDR_ < 0.001), High terrain vs. Low (both p_FDR_ < 0.05), Med terrain vs. Flat (both p_FDR_ = 0.044), and additional lower beta power in Med terrain vs. Low (p_FDR_ = 0.040) at the left sensorimotor cluster. At <u>the left posterior parietal cluster</u>, both age groups exhibited lower beta power reduction at greater terrain unevenness in all pairwise comparisons (all p_FDR_ < 0.005) except for High terrain versus Med. At the <u>mid-</u> <u>cingulate cluster</u>, both age groups also exhibited lower beta power reduction at greater terrain unevenness in all pairwise comparisons (all p_FDR_ < 0.01) except for Med terrain versus Low. At the <u>left pre-supplementary cluster</u>, both younger and older adults exhibited significant beta reduction at High terrain vs. Flat (p_FDR_ < 0.001), High terrain vs. Low (p_FDR_ = 0.013), and Med terrain vs. Flat (p_FDR_ = 0.006). Lastly, at the <u>right premotor cluster</u>, both younger and older adults exhibited significant beta reduction at High terrain versus Flat (p_FDR_ < 0.001) and Med terrain vs. Flat (p_FDR_ = 0.002).

### Intra-stride ERSP

We computed the event-related spectral perturbations (ERSPs) tied to gait events and normalized to the average power at each frequency across the gait cycle for each condition. We also obtained the peak-to-peak ERSP for each band (theta, alpha, beta) for each terrain condition for younger and older adults (**Figure 8 – 10, Supplementary** Figure 7).

**Figure 8:**
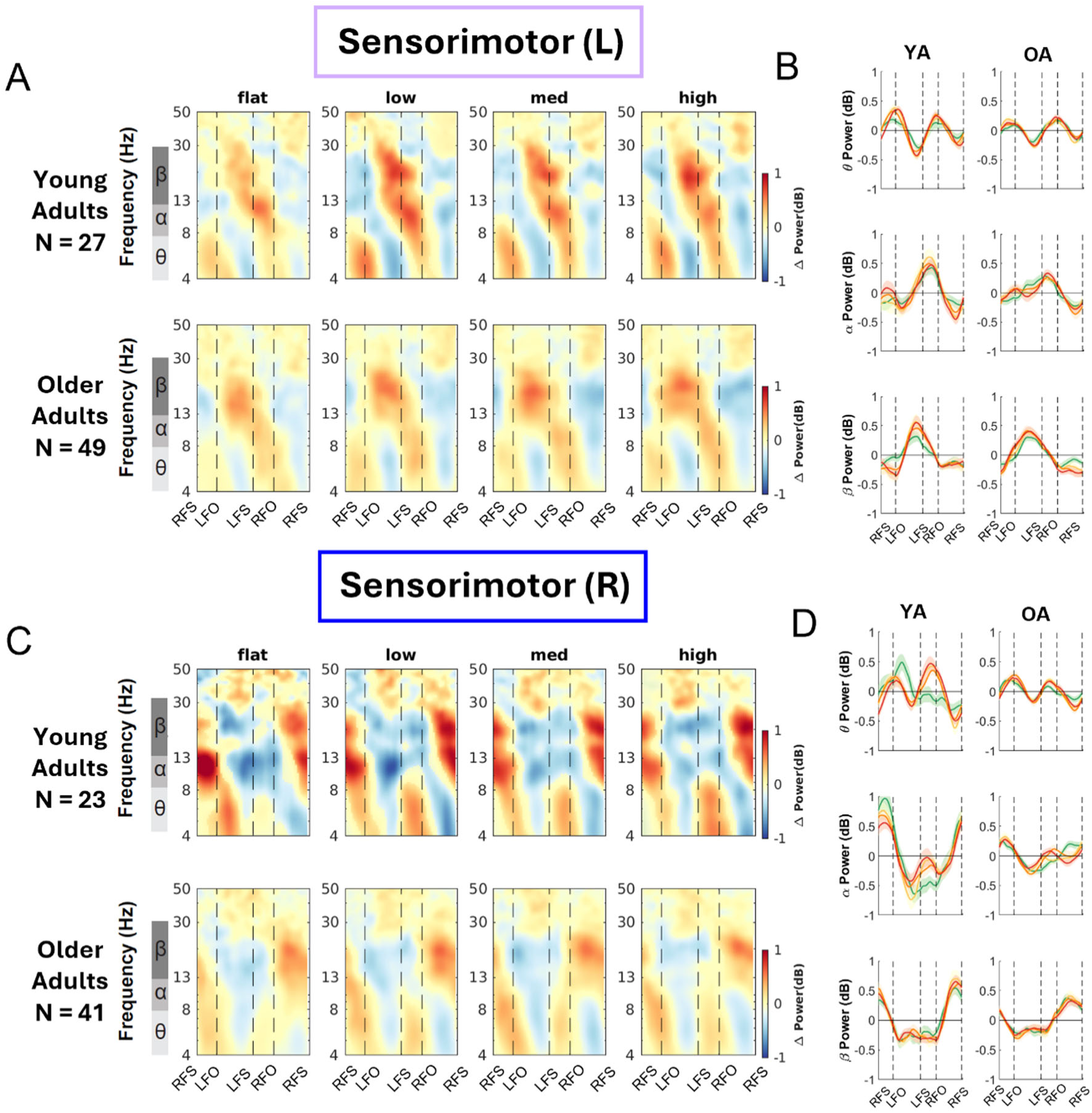
Intra-stride event-related spectral perturbations (ERSPs) with respect to the average of condition at different terrains for younger and older adults at left sensorimotor area (A) and right sensorimotor area (B), average intra-stride power fluctuations for each band at the left sensorimotor area (C) and right sensorimotor area (D). (A, C) The ERSPs were displayed with the gait cycle on the x-axis (RFS: right foot strike; LFO: left foot off; LFS: left foot strike; RFO: right foot off). All unmasked colors are statistically significant spectral power fluctuations relative to the mean power within the same condition. Colors indicate significant increases (red, synchronization) and decreases (blue, desynchronization) in spectral power from the average spectrum for all gait cycles to visualize intra-stride changes in the spectrograms. These data are significance masked (p < 0.05) through nonparametric bootstrapping with multiple comparison correction using false discovery rate. (B, D) Average spectral perturbations for each band (theta, alpha, beta) across the gait cycle for younger and older adults. YA: younger adults. OA: older adults.

<u>Sensorimotor cluster</u>: There was lateralization in the alpha and beta bands for left and right sensorimotor clusters for both younger and older adults (**Figure 8**). Younger adults demonstrated alpha and beta desynchronization during contralateral swing phase and the subsequent double support phase, while alpha and beta synchronization during the contralateral single limb stance phase and push-off. Older adults primarily demonstrated such synchronization and desynchronization in the beta band but not in the alpha band.

<u>Posterior Parietal cluster</u>: We also observed within-stride ERSPs modulation in bilateral posterior parietal areas (**Figure 9**). There was theta and alpha power desynchronization during the swing phase and synchronization during the double support phase for both younger and older adults. Qualitatively, as the terrain becomes uneven, we observed more desynchronization during the contralateral swing phase compared to Flat terrain. For older adults, we mainly observed theta band modulation with desynchronization during the mid-swing and synchronization during the double support phase.

**Figure 9:**
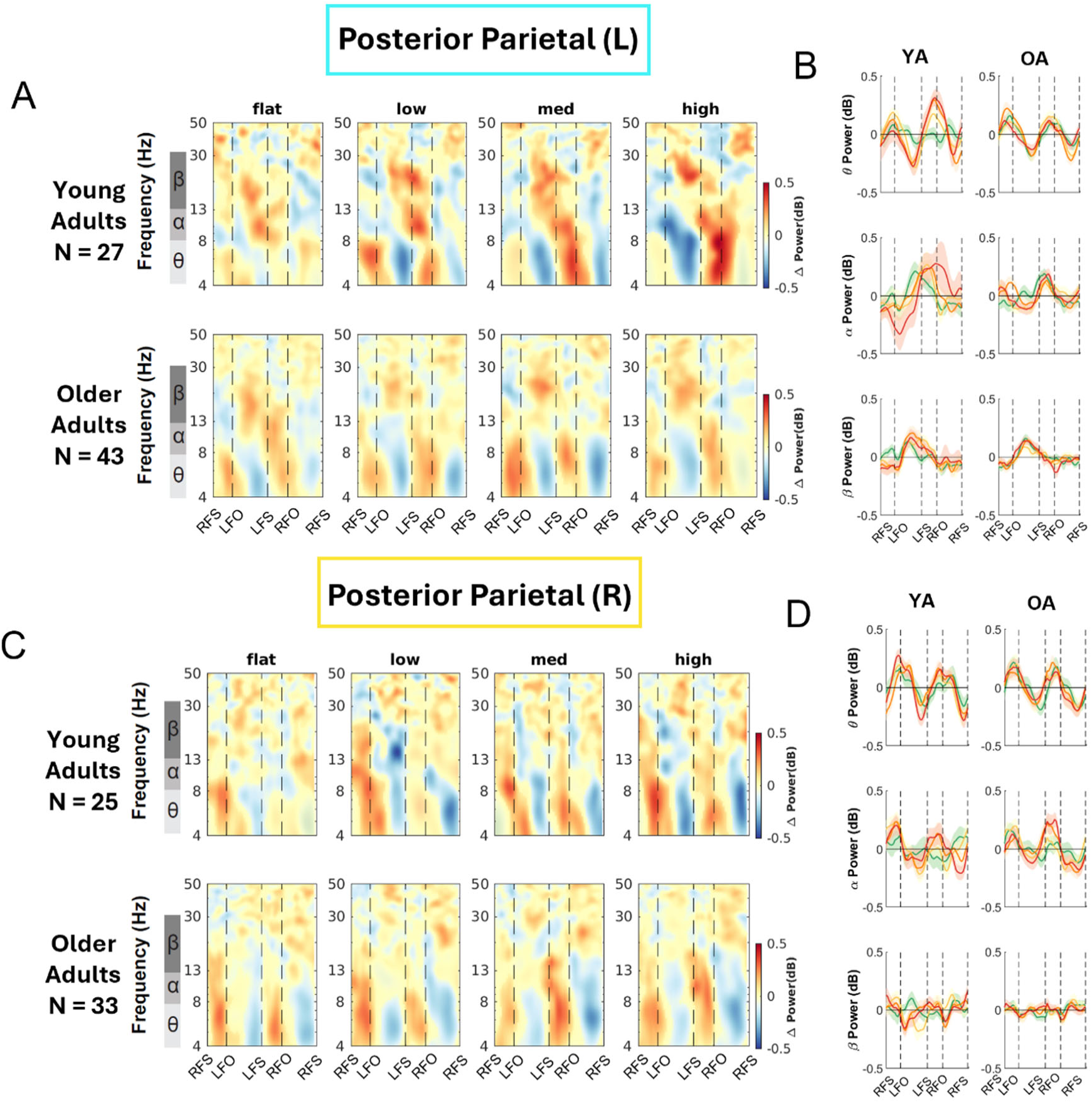
Intra-stride event-related spectral perturbations (ERSPs) with respect to the average of condition at different terrains for younger and older adults at left (A) and right posterior parietal area (B), average intra-stride power fluctuations for each band at the left (C) and right posterior parietal area (D). (A, C) The ERSPs were displayed with the gait cycle on the x-axis (RFS: right foot strike; LTO: left toe off; LFO: left foot off; RFO: right foot off). All unmasked colors are statistically significant spectral power fluctuations relative to the mean power within the same condition. Colors indicate significant increases (red, synchronization) and decreases (blue, desynchronization) in spectral power from the average spectrum for all gait cycles to visualize intra-stride changes in the spectrograms. These data are significance masked (p < 0.05) through nonparametric bootstrapping with multiple comparison corrections using false discovery rate. (B, D) Average spectral perturbations for each band (theta, alpha, beta) across the gait cycle for younger and older adults. YA: younger adults. OA: older adults.

Mid Cingulate cluster: At the mid cingulate area, we observed within-stride ERSPs modulation in mid cingulate area as theta and alpha power desynchronized during the swing phase and synchronized during the double support phase for both younger and older adults (**Figure 10AB**).

**Figure 10:**
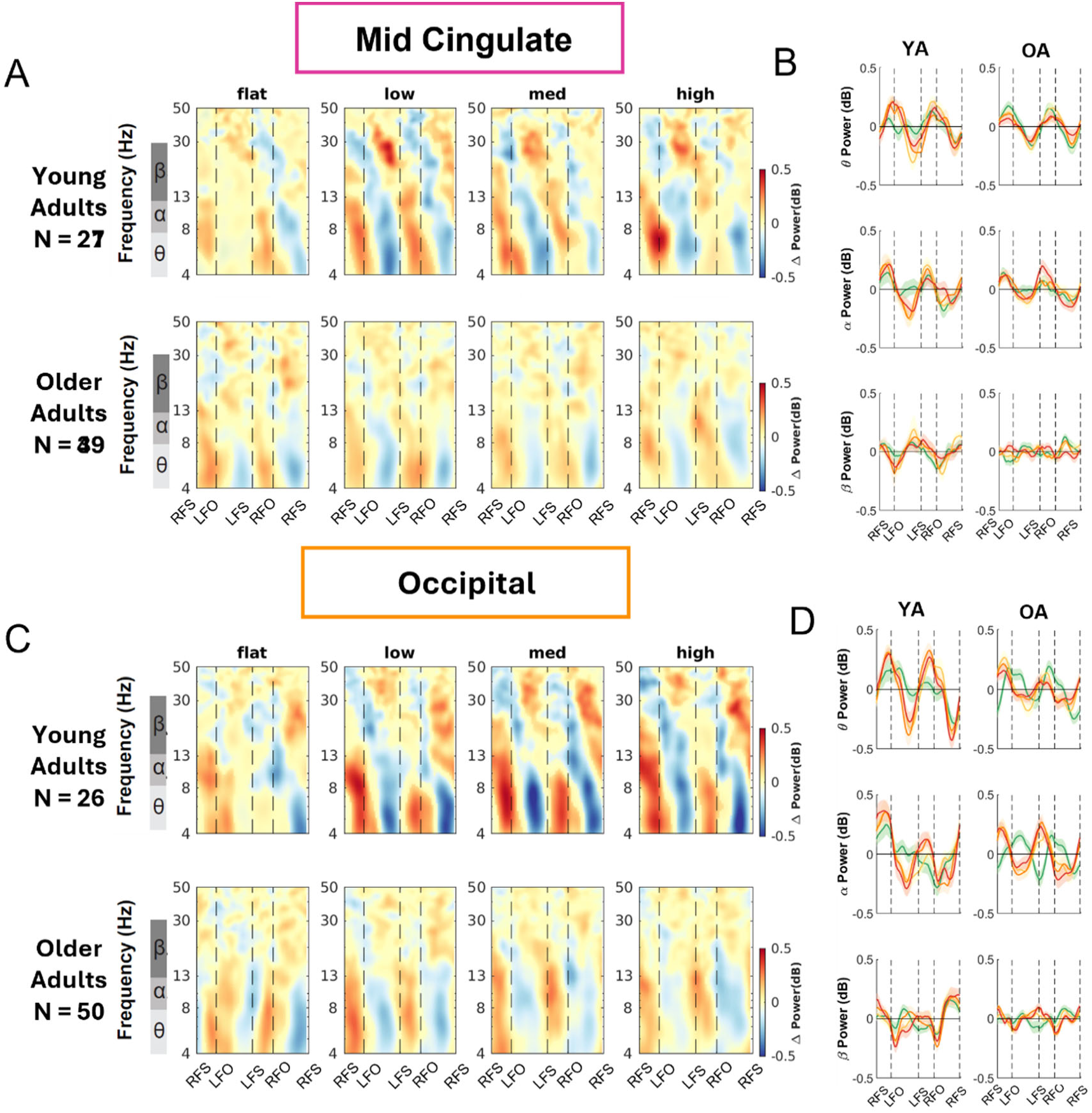
Intra-stride event-related spectral perturbations (ERSPs) with respect to the average of condition at different terrains for younger and older adults at mid cingulate (A) and occipital area (B), average intra-stride power fluctuations for each band at the mid cingulate (C) and occipital area (D). (A, C) The ERSPs were displayed with the gait cycle on the x-axis (RFS: right foot strike; LFO: left foot off; LFS: left foot strike; RFO: right foot off). All unmasked colors are statistically significant spectral power fluctuations relative to the mean power within the same condition. Colors indicate significant increases (red, synchronization) and decreases (blue, desynchronization) in spectral power from the average spectrum for all gait cycles to visualize intra-stride changes in the spectrograms. These data are significance masked (p < 0.05) through nonparametric bootstrapping with multiple comparison corrections using false discovery rate. (B, D) Average spectral perturbations for each band (theta, alpha, beta) across the gait cycle for younger and older adults. YA: younger adults. OA: older adults.

<u>Occipital cluster</u>: At the occipital area, we observed broadband (theta, alpha, and beta) synchronization during the double support phase and desynchronization around the mid-swing during uneven terrain walking for younger adults whereas the broadband spectral fluctuations were less prominent during Flat surface walking (**Figure 10CD**). For older adults, we also found significant broadband synchronization mainly during the double support phase and desynchronization during the midswing phase.

### Age and terrain unevenness on peak-to-peak ERSP

We tested whether theta intra-stride power fluctuations, alpha power fluctuations, and beta power fluctuations were affected by age, terrain, and interaction for each cluster after controlling for treadmill walking speed (**Figure 11**). Full statistical analysis results are reported in **Supplementary Table 13-15**.

**Figure 11.**
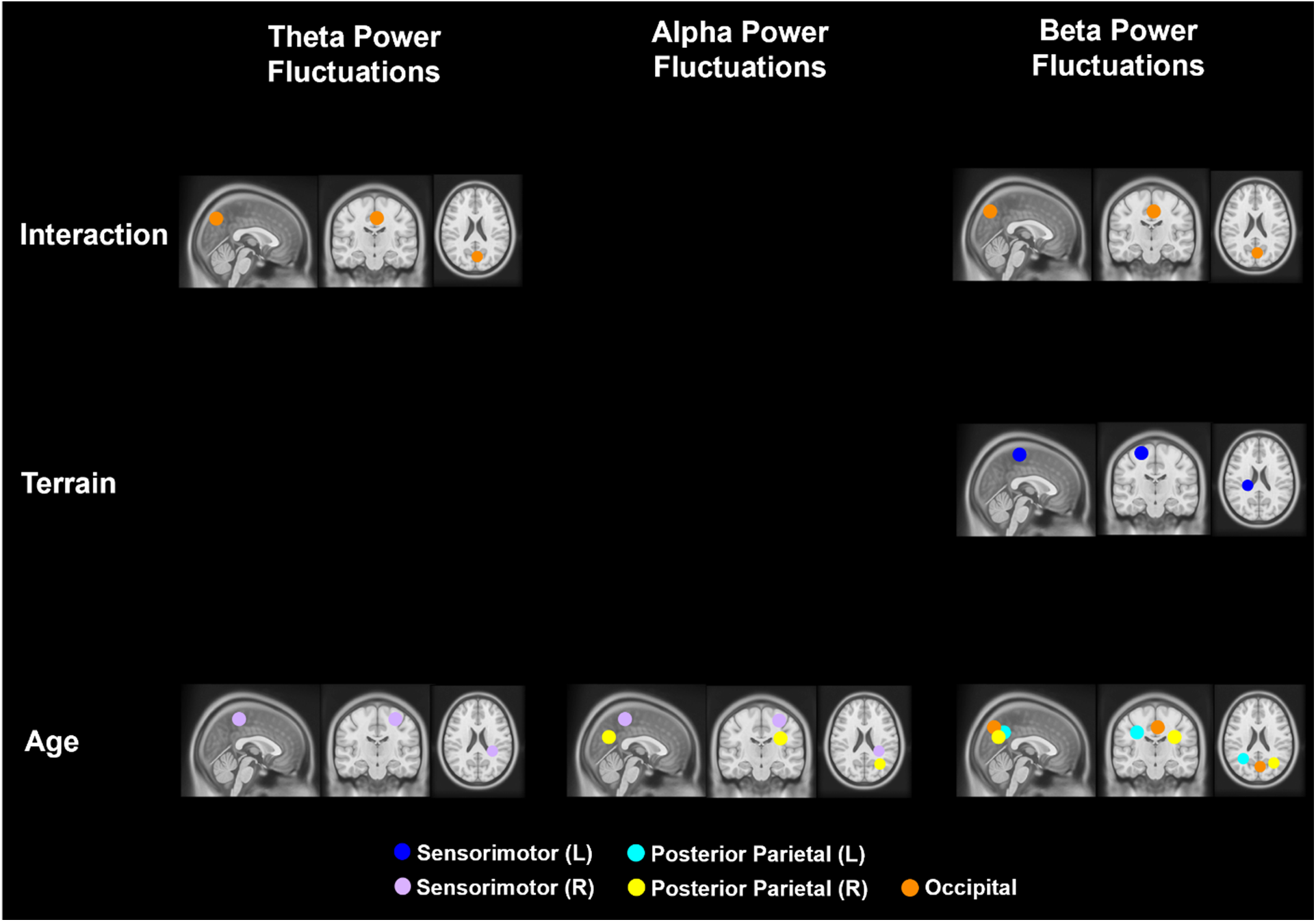
Significant effects of age, terrain, and interaction on theta, alpha, and beta band intra-stride power fluctuations (p < 0.05). The analyses were controlled for the effect of treadmill walking speed. Only the centroids of each cluster were plotted.

**Interaction effect:** The effect of terrain on EEG power fluctuations depends on age group primarily in the occipital cluster (**Figure 12A**). Significant interaction effect of age and terrain on peak-to-peak **theta band power fluctuations** was found in the occipital (F(3, 221) = 4.8, p = 0.003, η²ᵖ = 0.061) after controlling for treadmill walking speed.

**Figure 12:**
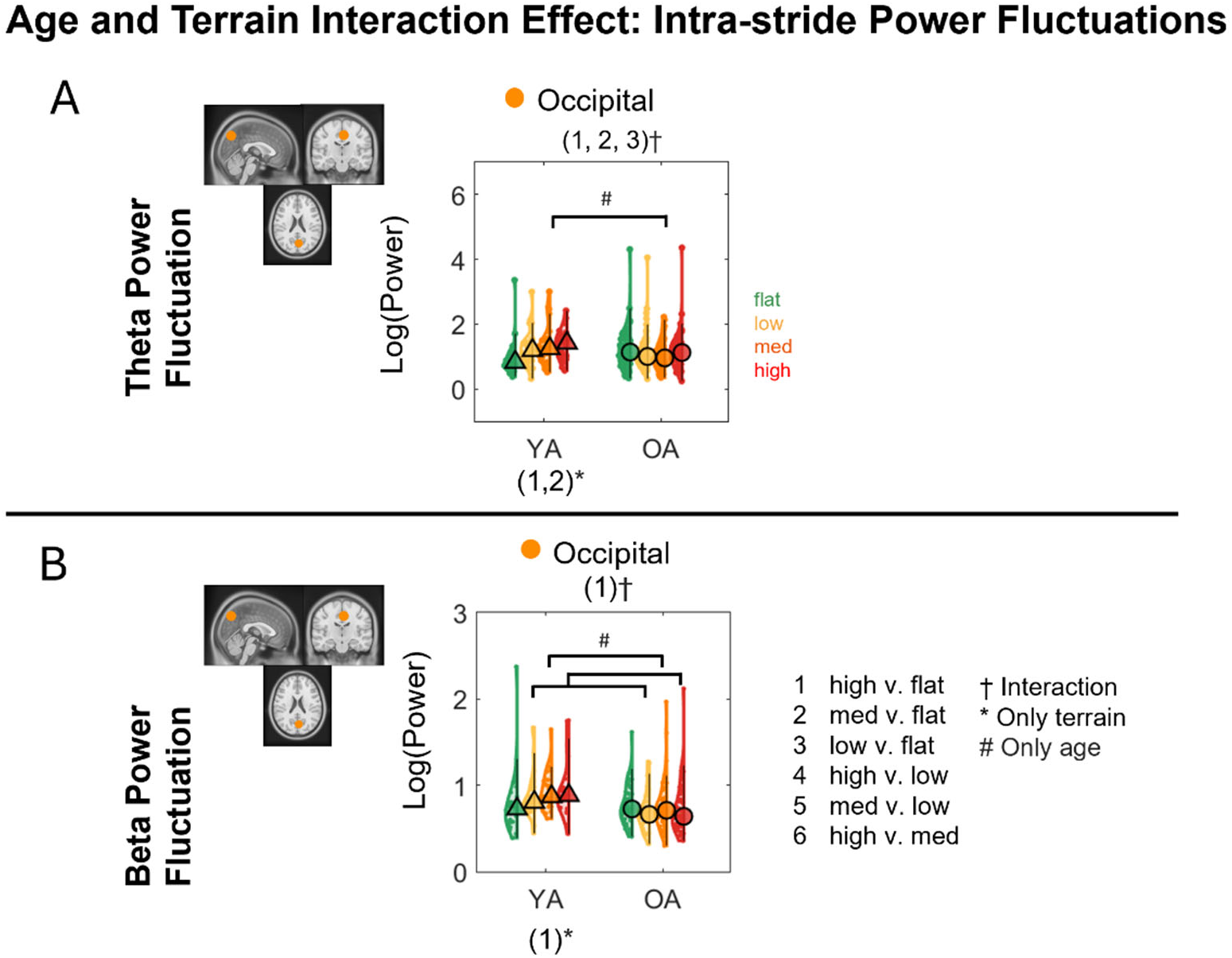
Violin plots of the average theta power fluctuations (A) and beta power fluctuations (B) for each terrain and age group for the corresponding brain clusters with a significant interaction effect. YA: younger adults, OA: older adults. #Indicates significant group differences between two terrain conditions, *Indicates significant differences (p < 0.05) between terrain conditions. †Indicates significant interaction effects between terrain and age group. 1: High vs. Flat, 2: Med vs. Flat, 3: Low vs. Flat, 4: High vs. Low, 5: Med vs. Low, 6: High vs. Med. Circle and triangle marker indicate median across participants.

Compared to younger adults, older adults had less power fluctuation in High terrain vs. Flat (p_FDR_ = 0.016), Med terrain vs. Flat (p_FDR_ = 0.003), and Low terrain vs. Flat (p_FDR_ = 0.047). Additionally, younger adults demonstrated greater power fluctuations at greater terrain unevenness at High versus Flat (p_FDR_ = 0.007) and Med versus Flat (p_FDR_ = 0.009). Theta power fluctuation was greater in younger adults than older adults at Med terrain (p_FDR_ = 0.027).

A significant interaction effect of age and terrain on **beta power fluctuations** was found in the <u>occipital</u> cluster (F(3, 221) = 3.24, p = 0.023, η²ᵖ = 0.042) with a significant main effect of age (F(3, 73) = 13.36, p < 0.001, η²ᵖ = 0.16; **Figure 12B**) after accounting for walking speed. Compared to younger adults, older adults had less difference in power fluctuation in High terrain vs. Flat (p_FDR_ = 0.021). Additionally, younger adults had less power fluctuation in High terrain vs. Flat (p_FDR_ = 0.028) while older adults did not exhibit any changes in power fluctuation between terrains. Younger adults also had greater power fluctuations than older adults in High, Med, and Low terrain (all p_FDR_ < 0.01).

**Age effect.** There was a significant age effect on theta power fluctuations at the right sensorimotor cluster (F(1, 61) = 4.24, p = 0.044, η²ᵖ = 0.065), with greater alpha power fluctuations in younger adults than older adults (**Supplementary** Figure 8A). We also found a significant age effect on alpha power fluctuations at the right sensorimotor cluster (F(1, 61) = 16.2, p < 0.001, η²ᵖ = 0.21) and right posterior parietal cluster (F(1, 55) = 4.77, p = 0.033, η²ᵖ = 0.08), with greater alpha power fluctuations in younger adults than older adults (**Supplementary** Figure 8B). There was also a significant age effect on beta power fluctuations at the left posterior parietal cluster (F(1, 67) = 4.97, p = 0.029, η²ᵖ = 0.069) and right posterior parietal cluster (F(1, 55) = 12.18, p = 0.001, η²ᵖ = 0.181) with greater beta power fluctuations in younger adults than older adults (**Supplementary** Figure 8C).

**Terrain effect.** Only left sensorimotor cluster had a significant terrain effect on beta power fluctuations (F(3, 225) = 5.78, p = 0.001, η²ᵖ = 0.072). Pooled groups showed greater beta power fluctuations at High terrain vs. Flat (p_FDR_ = 0.004), Med terrain vs. Flat (p_FDR_ = 0.002), and Low terrain vs. Flat (p_FDR_ = 0.013; **Figure 13)**.

**Figure 13:**
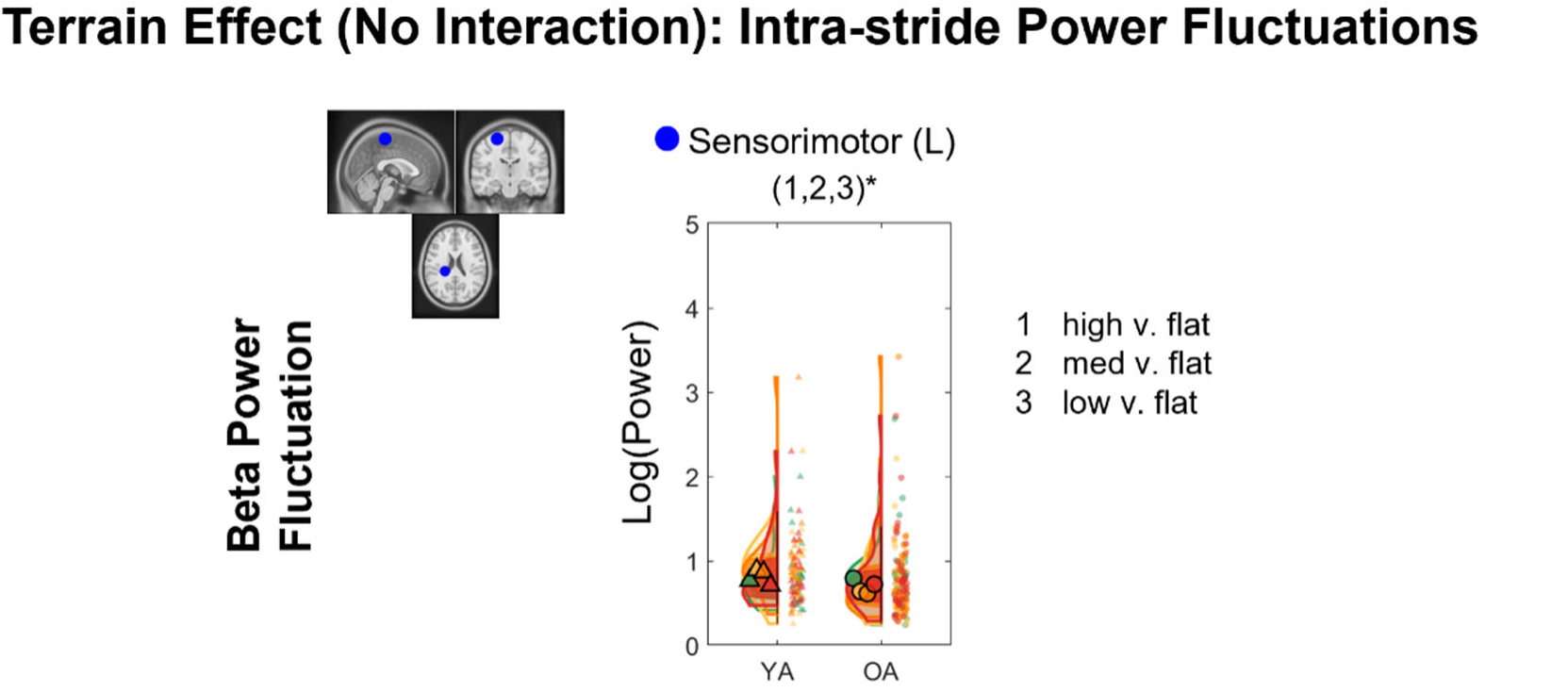
Violin plot of the beta power fluctuations with terrain unevenness at the left sensorimotor cluster. YA: younger adults, OA: older adults.

### Age and terrain unevenness on ERSP Sensorimotor Area

We found significant differences in ERSP using bootstrapping between uneven terrain and flat terrain in almost all brain areas (**Figure 14, Supplementary** Figure 9). We only found significant age-related differences in spectral fluctuations during uneven terrain walking in the left sensorimotor area, left posterior parietal, and occipital area (**Figure 14**). Red color indicates neural synchronization and blue color indicates neural desynchronization.

**Figure 14:**
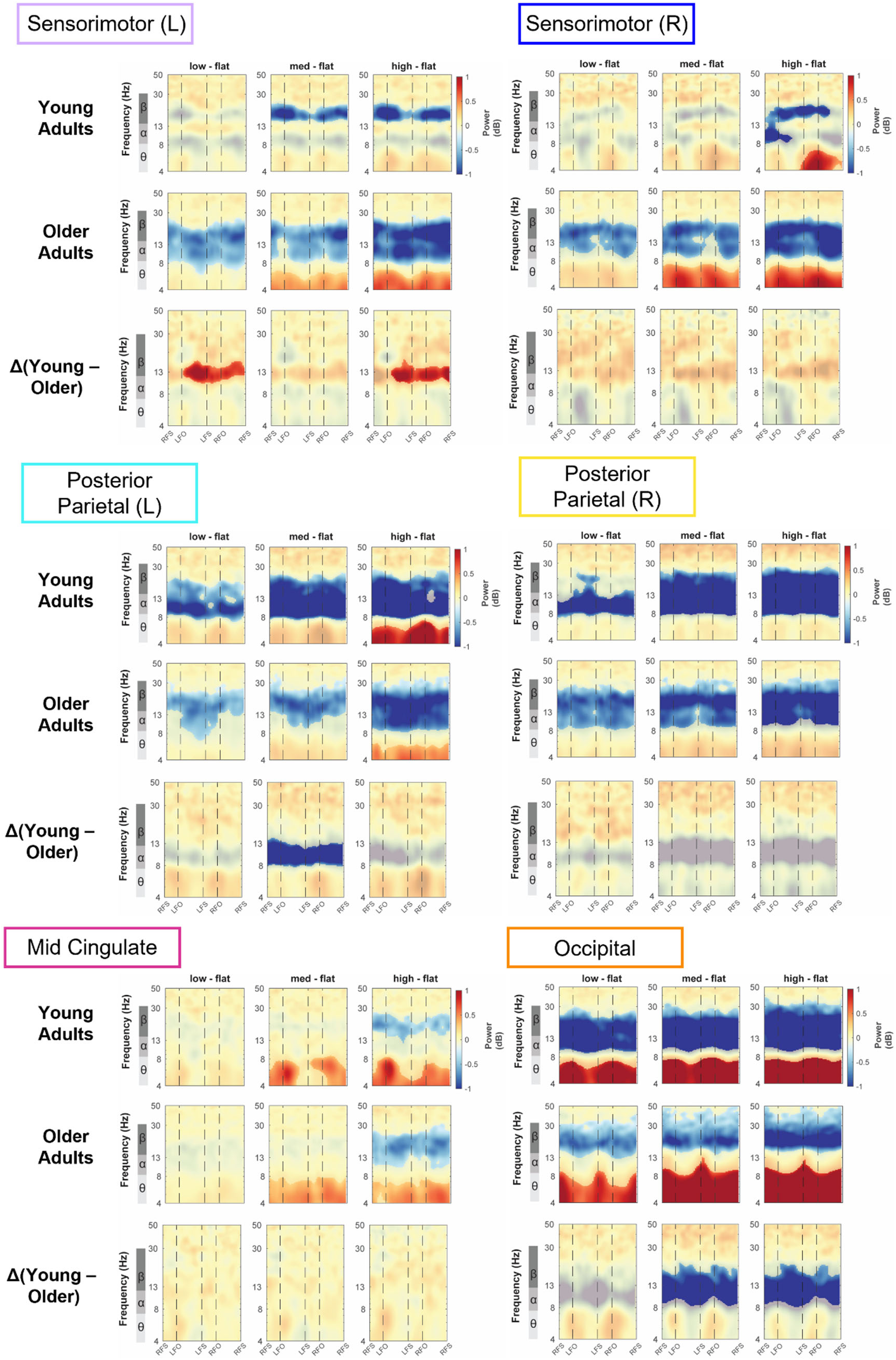
Average event-related spectral perturbations with respect to the Flat condition for each group and the difference between groups in the bilateral sensorimotor area, bilateral posterior parietal, mid cingulate, and occipital area. All unmasked colors are statistically significant spectral power fluctuations relative to the mean power within the same condition. Colors indicate significant increases (red, synchronization) and decreases (blue, desynchronization) in spectral power from the average spectrum for all gait cycles. The x-axes of the ERSPs are time in the gait cycle (RFS: right foot strike; LFO: left foot off; LFS: left foot strike; RFO: right foot off).

We observed significant spectral fluctuations in the sensorimotor areas both within age groups and between groups (**Figure 14**). With respect to Flat terrain, younger adults showed a significant increase in theta power during the double support phase during the High terrain condition in the right sensorimotor area, and older adults showed an increase in theta power in all uneven terrain conditions. Younger adults showed beta desynchronization during Med and High terrain primarily during the contralateral swing phase while older adults demonstrated alpha and beta desynchronization throughout the gait cycle during all uneven terrain surfaces in both left and right sensorimotor areas. Younger adults showed less alpha desynchronization from the contralateral single support phase to the contralateral swing phase for all uneven terrains than older adults, but only during the Low and High terrain were there significant differences between groups. There was no age group difference in the theta band.

### Posterior Parietal Area

There were significant spectral fluctuations in the posterior parietal areas both within age groups and between groups (**Figure 14**). With respect to Flat terrain, younger adults showed alpha desynchronization in Low, Med, and High terrain conditions and beta desynchronization in Med and High terrain conditions. Younger adults also exhibited theta synchronization during High terrain condition only at the left posterior parietal area. On the other hand, older adults demonstrated strong beta desynchronization throughout the gait cycle during all uneven terrain surfaces and a strong alpha desynchronization during the High terrain condition. When comparing older and younger adults, younger adults showed greater alpha desynchronization throughout the gait cycle during Med and High terrains, but only during Med terrain did it reach significance in the left posterior parietal area.

### Mid Cingulate

The mid-cingulate area demonstrated only a significant effect of terrain on spectral fluctuations (**Figure 14**). Both younger and older adults showed significant theta synchronization during Med and High terrain compared to Flat terrain, with additional beta desynchronization during High terrain. There was no evident difference between younger and older adults.

### Occipital Area

There were significant spectral fluctuations in the occipital area both within age groups and between groups (**Figure 14**). With respect to Flat terrain, younger adults showed theta synchronization and alpha/beta desynchronization throughout the gait cycle in Low, Med, and High terrain conditions. Older adults demonstrated theta synchronization and beta desynchronization throughout the gait cycle during all uneven terrain surfaces. They also exhibited alpha desynchronization during the double support phase during the medium and high terrain conditions. When comparing older and younger adults, younger adults showed greater alpha and lower beta band desynchronization throughout the gait cycle during Med and High terrain.

### Pre-supplementary and premotor area

For left pre-supplementary and right premotor area, we observed significant spectral fluctuations within age groups only (**Supplementary** Figure 9). At the left pre-supplementary area and right premotor area, younger adults showed theta synchronization during the double support phase and beta desynchronization at High terrain conditions compared to Flat terrain. Older adults only showed theta synchronization at Med and High terrain at the left pre-supplementary area. There was no evident difference across terrain unevenness.

## Discussion

This study’s primary objective was to determine whether there are age differences in electrocortical activity measured by EEG during walking on parametrically varied uneven terrain. Our results showed that, compared to younger adults, older adults walked slower on uneven terrain and had greater increases in step duration variability and mediolateral sacral excursion variability with terrain unevenness. Contrary to our hypothesis, younger adults experienced a greater reduction in alpha and beta power in the parietal-occipital region compared to older adults when terrain unevenness increased. Additionally, our results demonstrated that walking on uneven terrain leads to widespread changes in electrocortical dynamics in the brain, especially in alpha and beta band power. We also assessed how intra-stride power fluctuations changed with terrain unevenness and age. Gait-related fluctuations in the occipital theta and beta bands increased with terrain unevenness in younger adults but not in older adults. Older adults had smaller intra-stride theta and alpha band power fluctuations than younger adults in the right sensorimotor area and smaller intra-stride beta band power fluctuations at the posterior parietal areas.

Gait-related spectral perturbations also revealed several age differences in intra-stride power fluctuations. The greatest age difference was in the occipital area, where older adults showed less alpha and beta desynchronization within walking strides on an uneven surface compared to on a flat surface. At the left sensorimotor area, alpha desynchronization at Low terrain and High unevenness was greater in older adults than younger adults, while at the left posterior parietal area, beta desynchronization at Med terrain unevenness versus Flat was less in older adults. In summary, older adults showed a greater increase in gait variability with greater terrain unevenness than younger adults but exhibited a lack of modulation of parieto-occipital activity in response to terrain unevenness. This lack of neural response may reflect a reduced capacity to enhance visuomotor processing on uneven terrain, potentially resulting in poorer gait performance.

It is worth noting that since older adults walked slower than younger adults, speed may have affected the EEG spectral measures and masked age-related changes. To disentangle the effect of walking speed on EEG power, our recent paper used the same cohort of participants to examine the effect of walking speed (0.25m/s to 1.0m/s) and age on EEG spectral power. First, older adults only showed a greater increase in left posterior parietal theta band power with speed than younger adults among all brain areas. This brain area and band do not overlap with our main finding that in the parieto-occipital region, alpha and beta band power showed the largest age differences in response to changes in terrain unevenness. Additionally, there is widespread alpha and beta band power reduction with faster walking speed in both younger and older adults. The reduction of EEG power was about 0.071 dB for a walking speed of 0.39 m/s (average walking speed for older adults) to 0.70 m/s (average walking speed for younger adults). However, the age-related differences in parieto-occipital EEG power between terrain conditions are at least 10-20 times larger than the effect of walking speed. Therefore, it is unlikely that walking speed is the main contributor to changes in EEG power spectral densities.

### Widespread effect of terrain on theta, alpha, and beta band power

Theta spectral power slightly increased with terrain unevenness for both younger and older adults at the mid-cingulate and posterior parietal areas (mostly between the High terrain vs Flat terrain) but sharply increased with terrain unevenness at the occipital area (**Figure 7**). The observed mid-cingulate theta band activity is consistent with the evidence that midfrontal theta activity is associated with error detection and monitoring (Cohen, 2014; Ficarella et al., 2019). The occipital theta band power increases with terrain unevenness. Our results may be explained by the fact that greater occipital theta band power is associated with greater heterogeneity in the whole visual field (Feldmann-Wüstefeld et al., 2017). Since the rigid pucks at greater terrain unevenness were more variable or heterogeneous in height, this could contribute to the observed increase in theta band power. In contrast, Yokoyama et al. found an opposite direction of change in theta band power in that participants reduced theta band activity when instructed to place their foot on visual targets during walking compared to normal walking (Yokoyama et al., 2021). The discrepancies between our results and that reported in Yokoyama et al. suggest that the rigid pucks on the treadmill may serve as distractors since they are not designed to be targets for foot placement. Future studies may be designed to investigate occipital theta band activity by dissociating distractors in the visual field and voluntary choice of stepping targets.

Alpha and beta spectral power decreased when walking on uneven terrains versus a flat surface for both younger and older adults in widespread brain areas, including sensorimotor, mid-cingulate, posterior parietal, and occipital areas (**Figure 7BC and Figure 14**). During the resting condition, alpha and beta oscillations are thought to be in the ‘idling’ state (Pfurtscheller et al., 1996a, 1996b). At the sensorimotor area, the reduction in beta power indicates greater cortical involvement for motor execution during uneven terrain walking compared to walking on a flat surface. These findings in the sensorimotor region are consistent with prior research that found beta band power reduction during more balance-challenging walking tasks, such as walking on a narrow beam (Sipp et al., 2013) and following immediate exposure to split-belt walking (N. A. Jacobsen & Ferris, 2023). Greater cortical involvement during balance-challenging walking tasks may be related to muscle co-contraction to increase joint stiffness for stability. Although there is no study that has directly investigated such a relationship, prior studies support the possibility. Greater cortico-muscular coherence in beta bands was observed when balance was perturbed for both younger and older adults compared to pre-perturbation quiet standing (Ozdemir et al., 2018), and muscle co-contraction became greater during dual-task walking, a paradigm with a higher cognitive load (Lo et al., 2017). Together, these findings suggest a potential connection between muscle co-contraction and cortical beta-band activity, but future study is needed.

At the posterior parietal area, reduction in alpha and beta band power was prominent throughout the gait cycle during uneven terrain walking compared to walking on flat surfaces (**Figure 14**). The posterior parietal cortex is responsible for sensory integration and visuospatial processing to aid whole-body motor planning (Drew & Marigold, 2015; Nordin et al., 2019). Here, we found that older adults also demonstrated sustained alpha and beta desynchronization throughout the gait cycle when walking on uneven surfaces, although the alpha band desynchronization was smaller than that of younger adults. This sustained alpha and beta desynchronization suggests a greater need for cortical processing of multi-sensory integration and greater attention to maintain balance on uneven surfaces (Liu et al., 2024).

### Parieto-occipital region alpha and beta bands exhibited the largest age-related differences in response to changes in terrain unevenness

Parietal-occipital region alpha and beta bands exhibited age differences in response to changes in terrain unevenness. Older adults had less reduction in parieto-occipital region alpha and beta band power when walking onto uneven terrain versus flat surface, when comparing to younger adults (**Figure 6**). These results suggest that older adults had less of an increase in the posterior parietal and occipital involvement walking over uneven terrain from a flat surface. While this does not align with our initial hypothesis, there are several explanations for what we observe. First, our results may fit into the posterior-anterior shift in aging (PASA) model, which states that less posterior activity is associated with greater frontal activity as we age (Davis et al., 2008).

Although our results cannot directly infer such an association, as we did not identify a prefrontal cluster, a previous paper from our large Mind in Motion project using functional Near-Infrared Spectroscopy (fNIRS) at the prefrontal region reported that older adults had greater prefrontal activation than younger adults throughout all terrain conditions (Hwang et al., 2024). Therefore, future research may aim to investigate whether the PASA model can be generalized to the motor domain, especially during walking.

Second, it is possible that older adults had greater reliance on vision to guide their foot placement even when walking on a flat surface, while younger adults did not rely as much on vision during typical walking. This is congruent with the observation that at the occipital area, alpha desynchronization occurred throughout the gait cycle when walking on uneven terrain versus flat terrain for younger adults, but for older adults, alpha desynchronization was not as prominent (**Figure 6A and Figure 14**). Prior studies suggested that a reduction in occipital alpha power is associated with the neural processing of visual input, as occipital alpha power was reduced when participants walked in a light condition compared to dark (Cao et al., 2020). Additionally, alpha-band desynchronizations occur in the occipital area in response to visual optical flow when seated (Vilhelmsen et al., 2015). Because we did not observe a reduction in occipital alpha power in older adults while walking on uneven surfaces, we speculate that older adults already rely heavily on visual input during walking on flat surfaces. If older adults engage parieto-occipital activity more than younger adults when walking on flat surfaces, there would be less range available for modulation as terrain unevenness increases. In contrast, younger adults may only start to rely on visual input when walking on uneven surfaces.

In contrast to our hypothesis, we did not find age differences in sensorimotor power when walking on uneven terrains versus a flat surface. Previous work found greater beta power desynchronization in older adults than younger adults during wrist flexion/extension movement using the non-dominant hand (Espenhahn et al., 2019) and during button pressing (Bardouille & Bailey, 2019), although only a very small variance was explained (R^2^ = 0.064) between event-related beta desynchronization at motor cortex and age. Another study found that under a dual-task paradigm combining walking with a visual reaction time task, older adults had smaller beta power desynchronization change from sitting to walking compared to younger adults (Protzak & Gramann, 2021). One potential reason why we did not observe age differences in the sensorimotor area could be that we had participants walk at their self-selected walking speed across varying terrain conditions and controlled for walking speed in our statistical analysis.

Since older adults walked slower than younger adults, speed may have affected the sensorimotor area and masked age-related changes (Alcock et al., 2023), although our recent analysis also did not find age differences in sensorimotor activity modulated by walking speed using the same cohort of participants (Salminen et al., 2025).

### Intra-stride spectral power fluctuations change with terrain and age group

While the overall band power change may reflect a sustained movement-related neuronal state during many gait cycles, power fluctuations within the gait cycle may represent a gait phase-dependent local neuronal population activity associated with sensorimotor processing and integration (Seeber et al., 2014). Both groups of participants demonstrated an increase in intra-stride power fluctuations in the sensorimotor beta band (**Figure 13**). The event-related synchronization during the contralateral stance phase and desynchronization during contralateral leg swing increased with terrain unevenness. This may be because stepping on taller pucks alters foot pressure, leading to increased sensory feedback from the sole of the foot.

Additionally, the occipital area demonstrated the largest age differences in power fluctuations (**Figure 12**). Spectral fluctuations changed with terrain unevenness for younger adults at occipital theta and beta bands. Theta and beta fluctuations increased with terrain unevenness in younger adults but not in older adults. One possibility is that the lack of task-related power modulation may indicate reduced cortical network flexibility in older adults, as they may struggle to adapt to walking on novel uneven surfaces (Darna et al., 2025). It is also likely that smaller intra-stride power fluctuations in older adults compared to younger adults in the occipital area are related to greater reliance on vision. These findings also suggest that older adults already heavily rely on vision during flat surface walking and thus might be unable to modulate brain activity to cope with the visuomotor processing demands of walking on uneven surfaces. Overall, our findings suggest that with greater terrain difficulty, increased intra-stride power fluctuations in sensorimotor regions may reflect a higher need for step adaptation, while those in the occipital area reflect greater reliance on visual information.

There appears to be a phase shift in gait-related power when comparing the flat condition and uneven terrains, especially for older adults (**Figure 10D**). When walking on uneven surfaces, the alpha-band showed synchronization during the double support and desynchronization during the single support phase. When walking on a flat surface, alpha synchronization occurred following foot off, while desynchronization occurred during double support. This result suggests that the visual processing timing may be different for flat versus uneven terrain walking for both younger and older adults. Future research should investigate the connectivity between the occipital area and sensorimotor region to delineate how the neural oscillations at different regions coordinate to facilitate walking.

Multiple motor-related regions showed greater power fluctuations in younger adults than older adults, including right sensorimotor theta and alpha-band fluctuations, right posterior parietal alpha-band fluctuations, and bilateral posterior parietal beta-band fluctuations, after adjusting for walking speed (**Supplementary** Figure 8). Combined with the slower walking speeds in our older adults, this finding is consistent with prior literature that demonstrated people with mobility deficits had reduced intra-stride fluctuations (Guo et al., 2024). For example, people with Parkinson’s Disease demonstrated reduced intra-stride power fluctuations in the alpha and beta bands compared to control participants (Guo et al., 2024). Since greater gait variability in stride length and stride time is associated with smaller alpha band fluctuations, it is possible that the smaller alpha power fluctuations in older adults also result from greater gait variability in older adults than in younger adults (Downey et al., 2022).

### Limitations

There are several limitations to our study, including the inherently relatively low spatial resolution of EEG compared to other imaging modalities such as fMRI and the lack of gaze data to fully understand visuospatial processing during uneven terrain walking (Liu et al., 2024). Interestingly, we found that older adults had fewer brain components compared to younger adults. Almost 10% of older adults do not have brain components or have <5 brain components, resulting in more participants having to be dropped from the analysis. Several factors may explain the age differences in the number of brain components. Dry or thickened scalp skin due to aging may increase the EEG impedance. Heavy breathing and neck or facial muscle activity due to task difficulty can lead to greater contamination of the brain signal. It is possible that ICLabel mislabeled brain components for older adults, as this toolbox was primarily trained on data from younger adults. Another explanation is that older adults demonstrate non-selective recruitment of brain regions relative to younger adults (Bunzeck et al., 2024; Seidler et al., 2010). For example, when performing motor tasks with the dominant hand, older adults activated both contralateral and ipsilateral brain regions during simple motor tasks, whereas younger adults show largely contralateral motor cortical activation (Riecker et al., 2006). It is possible that EEG source separation had difficulty with bilateral brain sources for older adults that had unilateral sources in younger adults, leading to a reduced number of brain components in the older adults.

Another limitation is the imbalance in sample sizes between young (n = 31) and older adults (n = 71) when forming brain clusters using K-means. Although our statistical analyses for power spectral densities and event-related spectral perturbations accounted for this unequal sample size, the K-means clustering used to form brain clusters may have been biased toward dipole locations from older adults. Consequently, the resulting cluster centroids likely reflected a greater influence from the older adult group. Future studies could address this by using a weighted K-means approach to account for unequal group sizes.

In the present study, we did not identify a cluster of brain sources in the prefrontal area, despite prior findings using functional near-infrared spectroscopy showing increased prefrontal activity in older adults during walking on more uneven terrain (Hwang et al., 2024). Prefrontal activity can be difficult to obtain with mobile EEG using ICA due to ocular artifacts and facial muscle activities. Additionally, ICA only recovers the strongest brain sources in EEG data averaged across many gait cycles. Since our study did not incorporate a cognitively intensive task during walking, prefrontal sources may not be sufficiently strong compared to the ongoing rhythmic electrocortical fluctuations dominant during steady state gait. If there was a series of arrhythmic and discrete prefrontal neural activations involved in gait adjustments to the uneven terrain, it would produce a robust fNIRS signal but a relatively weak electrocortical signature when viewed in comparison to the strong electrocortical rhythms that occur during steady state gait. This is reflective of the differences in neural metrics used by fNIRS and EEG (Richer et al., 2024).

## Conclusion

Walking on uneven terrain led to greater gait variability and widespread changes of electrocortical dynamics in the brain for both younger and older adults. We found the most prominent age differences in response to uneven terrain walking in the parieto-occipital region alpha and beta bands. Younger adults demonstrated a greater reduction in alpha and beta power with greater terrain unevenness, compared to older adults. Intra-stride power spectral fluctuations only changed with terrain unevenness for younger adults at the occipital area, but did not change with terrain unevenness for older adults. To summarize, older adults showed a greater increase in gait variability with greater terrain unevenness than younger adults but exhibited a lack of modulation of parieto-occipital activity in response to terrain unevenness. The lack of task-related EEG power modulation may indicate reduced cortical flexibility in older adults, resulting in poorer gait performance. These findings may also suggest that older adults already heavily rely on vision during flat surface walking and are thus unable to modulate brain activity to cope with the visuomotor processing demands of walking on uneven surfaces. Our future study will systematically investigate the association between behavioral changes and EEG spectral measures to determine whether this lack of modulation reflects neural compensation to maintain gait performance while walking on uneven terrain.

## Acknowledgement

We would like to thank Human Neuromechanics Lab members for their assistance with data collection: Madison Tenerowicz, Quinlan Degnan, Sydney Irwin, Sofia Arvelo Rojas, Tyler Irby, and Sai Shrestha, and other lab members for the insightful discussion that helped improve the paper. We would also like to thank our study coordinators for recruiting participants and maintaining the study database.

## Ethics Statement

The studies involving humans were approved by the Institutional Review Board (IRB) at the University of Florida (IRB 201802227). The studies were conducted in accordance with the local legislation and institutional requirements. The participants provided their written informed consent to participate in this study.

## Data Availability

Data for younger adults is available via OpenNeuro: https://openneuro.org/datasets/ds004625/versions/1.0.2.

Data for older adults is available for review via OpenNeuro: https://openneuro.org/datasets/ds006095

Code is available via GitHub Repo: https://github.com/changliu-99/MindInMotion_AgeDifferences_UnevenTerrain

## Author Contribution

**Chang Liu**: Methodology, Software, Formal analysis, Investigation, Data Curation, Writing – Original Draft, Visualization. **Erika M. Pliner**: Methodology, Investigation, Data Curation, Writing – Review & Editing. **Jacob S. Salminen**: Methodology, Software, Formal analysis, Investigation, Data Curation, Writing – Review & Editing. **Ryan J. Downey**: Methodology, Software, Formal analysis, Investigation, Data Curation, Writing – Review & Editing. **Jungyun Hwang**: Methodology, Investigation, Data Curation, Writing – Review & Editing. **Arkaprava Roy:** Formal analysis, Investigation, Writing – Review & Editing. **Ryland Swearinger**: Data curation, Investigation, Writing – Review & Editing. **Natalie Richer**: Methodology, Investigation, Data Curation, Writing – Review & Editing. **Chris J. Hass**: Conceptualization, Methodology, Investigation, Resources, Writing – Review & Editing, Project administration, Funding acquisition. **David J. Clark**: Conceptualization, Methodology, Investigation, Resources, Writing – Review & Editing, Project administration, Funding acquisition. **Todd M. Manini**: Conceptualization, Methodology, Investigation, Resources, Writing – Review & Editing, Project administration, Funding acquisition. **Yenisel Cruz-Almeida**: Conceptualization, Methodology, Investigation, Resources, Writing – Review & Editing, Funding acquisition. **Rachael D. Seidler**: Conceptualization, Methodology, Investigation, Resources, Writing – Review & Editing, Project administration, Funding acquisition. **Daniel P. Ferris**: Conceptualization, Methodology, Investigation, Resources, Writing – Review & Editing, Supervision, Project administration, Funding acquisition.

