## Supplementary Materials for "Age differences in electrocortical dynamics during uneven terrain walking"

### Supplementary Material

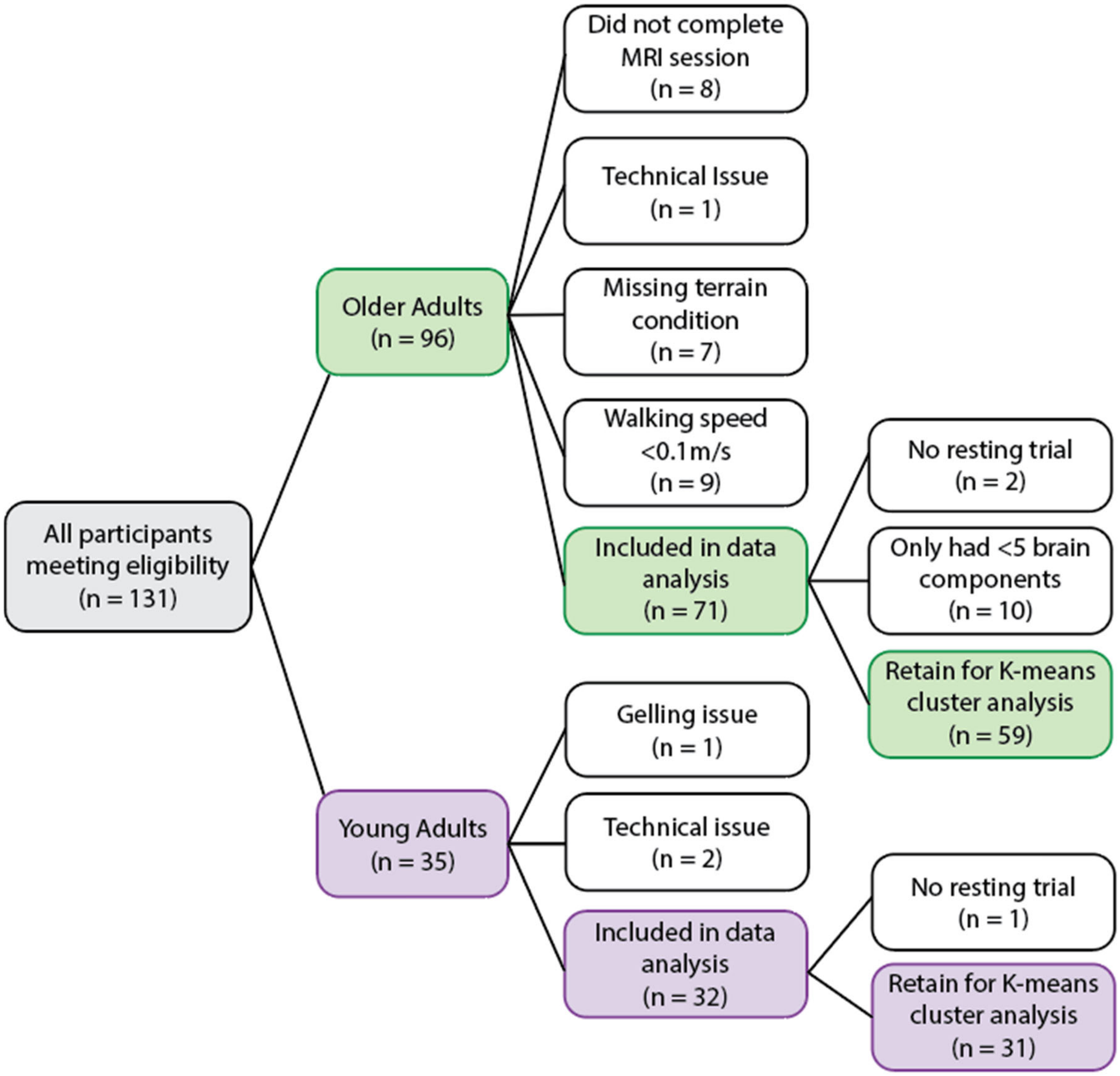


**Supplementary Figure 1**: Flow diagram of participant inclusion. We recruited a total of 35 young adults and 96 older adults for the EEG assessment. Among them, a total of 3 young adults were excluded for data analysis due to gelling issues with braids (n = 1) and 2) technical issues (n = 2). We further excluded 12 older adults and 1 young adult during data analysis. 25 older adults were excluded from data analysis due to 1) not completing the MRI session (n = 8), 2) a technical issue occurred during data collection (n = 1), 3) not completing high terrain conditions (n = 7), or 4) walking with a speed slower than 0.1m/s because these participants tended to walk toward the front of the treadmill, paused their walking, and then let the treadmill transport them toward the back.

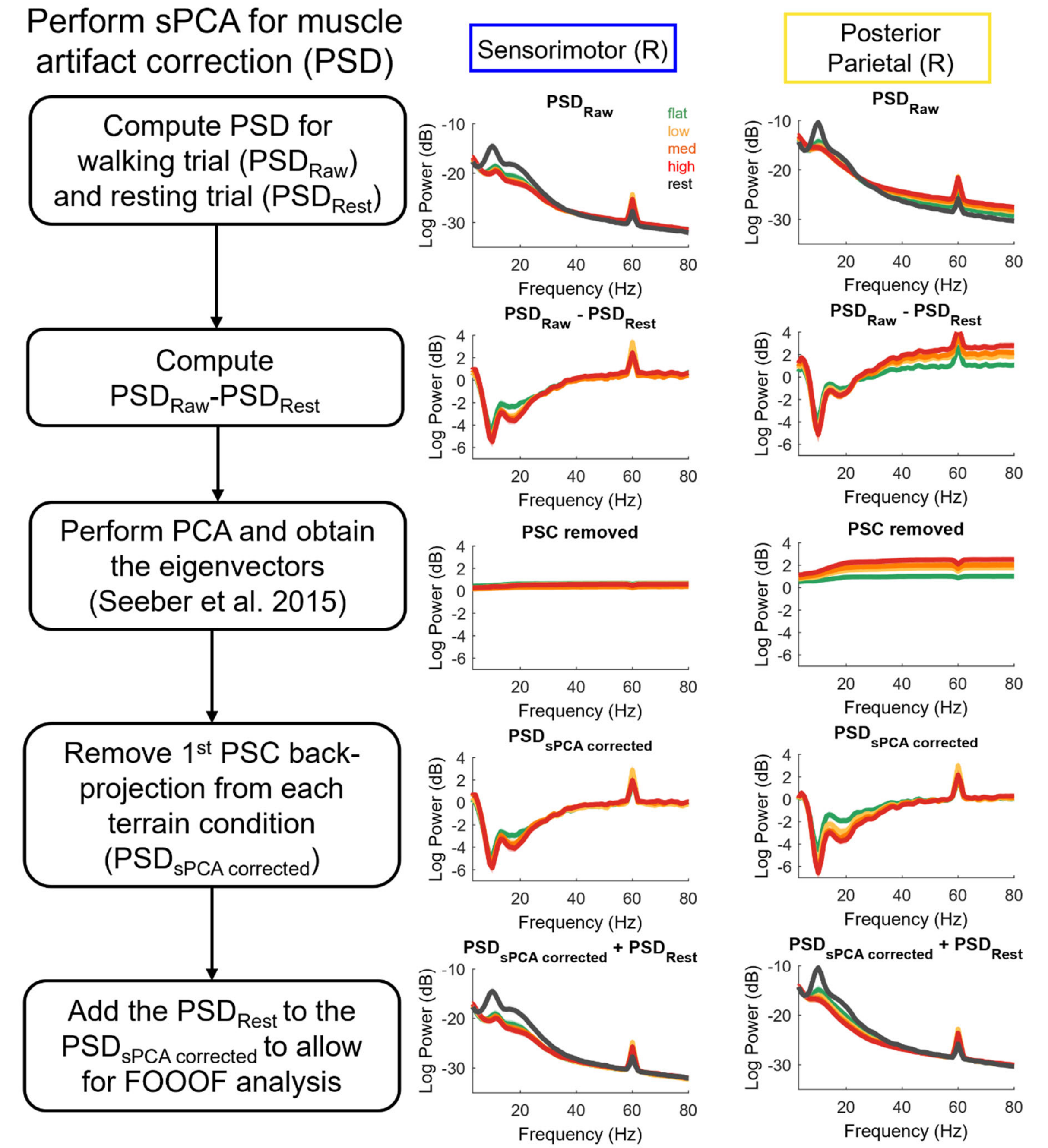


**Supplementary Figure 2:** Flow chart of spectral Principal Component Analysis for muscle artifact correction for the power spectral density analysis, along with the corresponding power spectral density plots at each step for the sensorimotor and posterior parietal areas. After obtaining the sPCA-corrected PSD, we added the PSD from the resting condition to the sPCA-corrected PSD to allow the FOOOF toolbox to separate the aperiodic and periodic components. PSC: principal component.

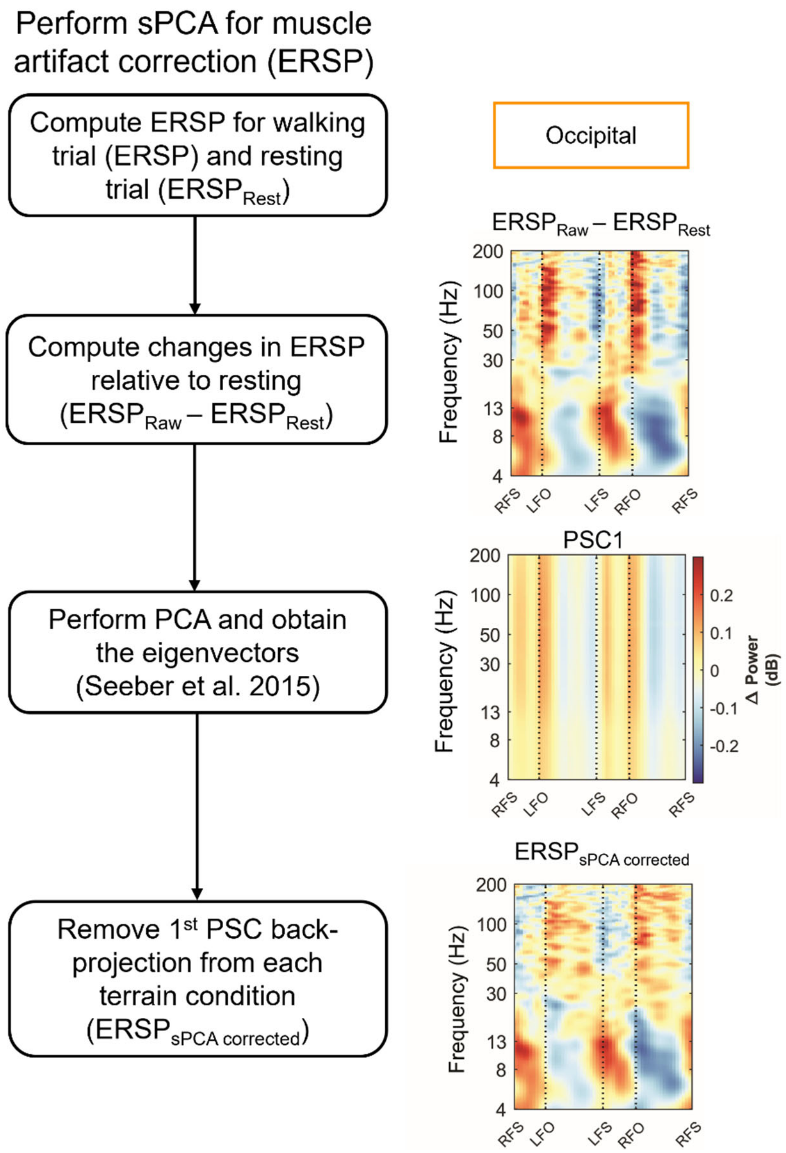


**Supplementary Figure 3:** Flow chart of spectral Principal Component Analysis for muscle artifact correction for the event-related spectral perturbation (ERSP), along with the corresponding ERSP plots at each step for the occipital cluster where the most muscle activity contamination occurred. PSC: principal component.

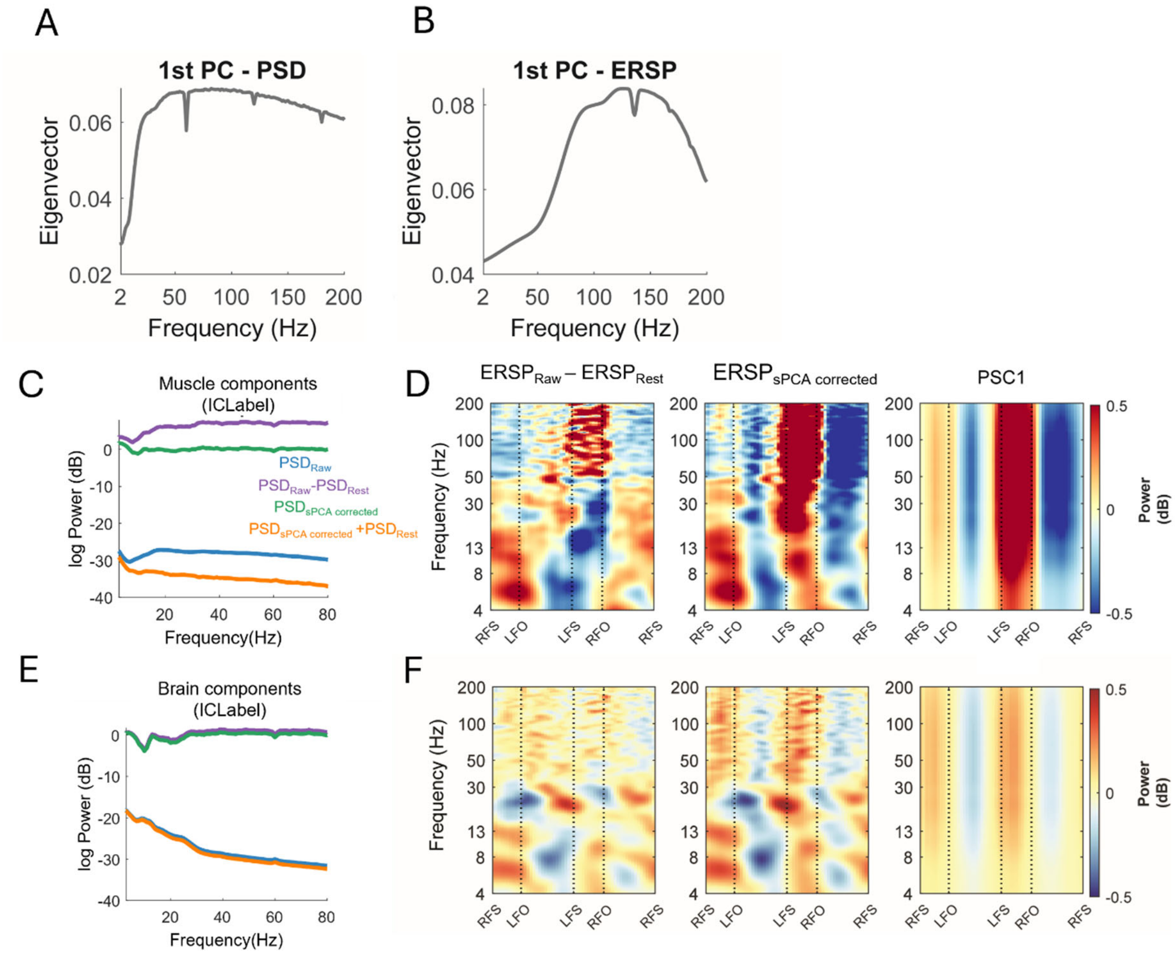


**Supplementary Figure 4:** Characteristics of the first spectral Principal Component. (A) First eigenvector for the power spectral densities (PSD; subject mean). (B) First eigenvector for the event-related spectral perturbations (ERSP; subject mean). (C) Averaged uncorrected and corrected PSDs for muscle components identified with ICLabel for one example participant. (D) Averaged uncorrected ERSP, sPCA corrected ERSP, and removed PSC1 for muscle components identified with ICLabel for one example participant. (E) Uncorrected and corrected PSDs for brain components identified with ICLabel for one example participant. (F) Uncorrected ERSP, sPCA corrected ERSP, and removed PSC1 for all brain components identified with ICLabel for one example participant.

Because the muscle components were corrected far more than the brain components, and the PSC1s mostly contained higher frequencies, we can reasonably conclude that PSC1 is primarily an electromuscular artifact.

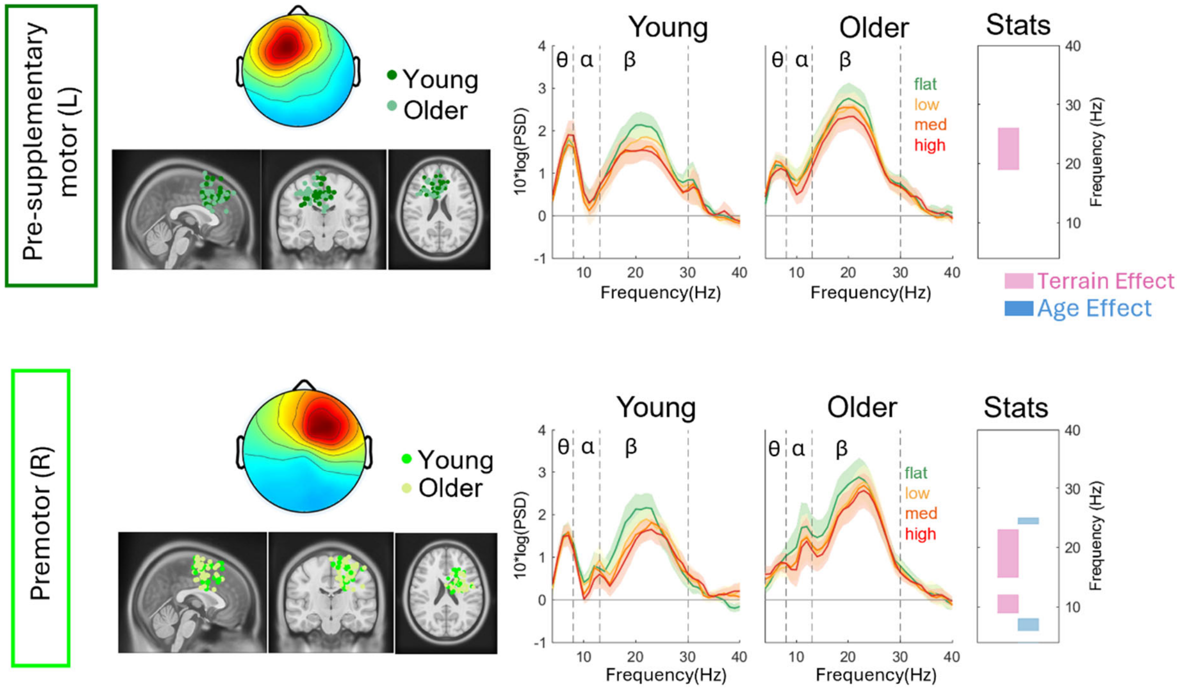


**Supplementary Figure 5**: Dipole location plotted on the Montreal Neurological Institute template, scalp topography, and average flattened power spectrum densities (PSDs) changed with terrain unevenness for younger and older adults for the pre-supplementary motor and premotor area. Shaded colored areas indicate standard error of PSDs across components in the cluster. Vertical black dashed lines indicate main frequency bands of interest—theta (4 - 8 Hz), alpha (8 - 13 Hz), and beta (13 - 30 Hz). The very right panel indicates the significant terrain and age effect on PSD at each frequency (pink: terrain, blue: age). In this non-parametric statistical analysis, only main effects were tested due to the limitation of this test.

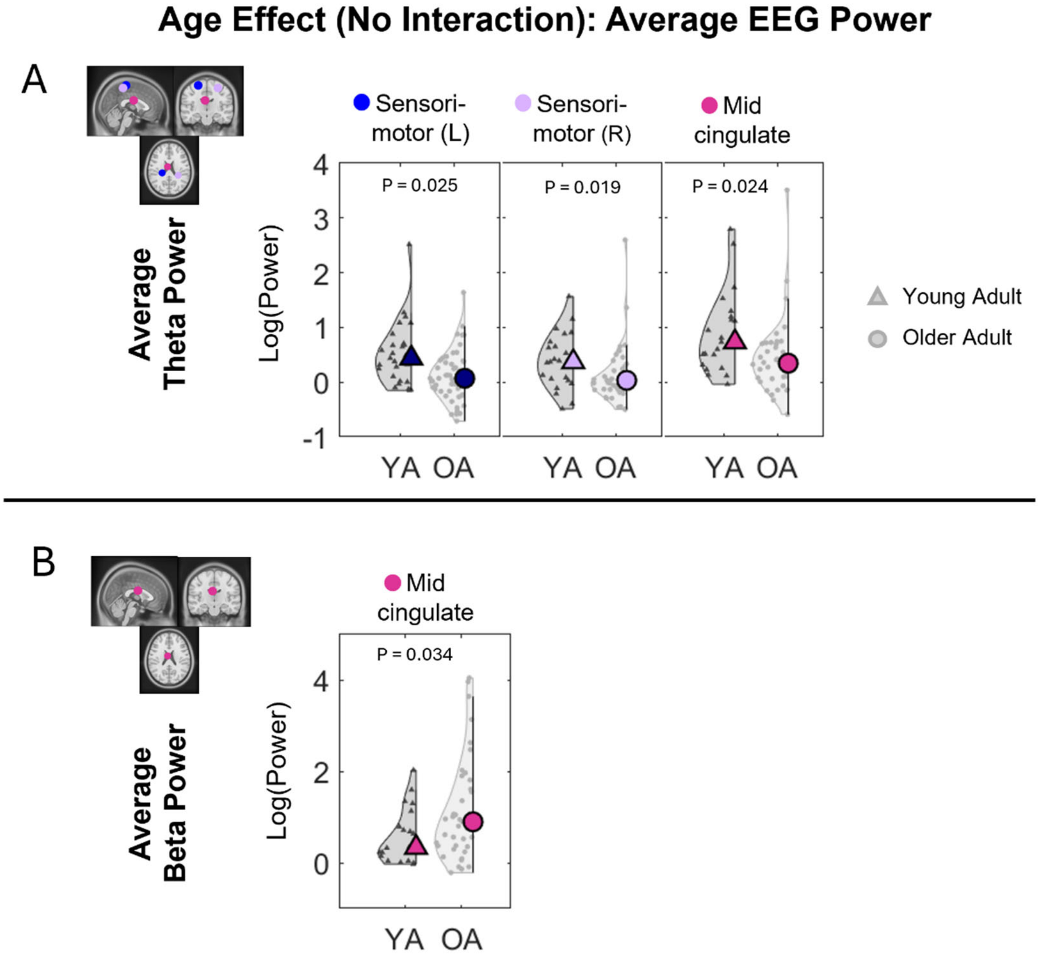


**Supplementary Figure 6**: Violin plots of the average alpha power (A) and beta power (B) for each age group after pooling all the terrain data for the corresponding brain clusters without an interaction effect. YA: younger adults, OA: older adults.  Circle and triangle markers indicate median across participants.

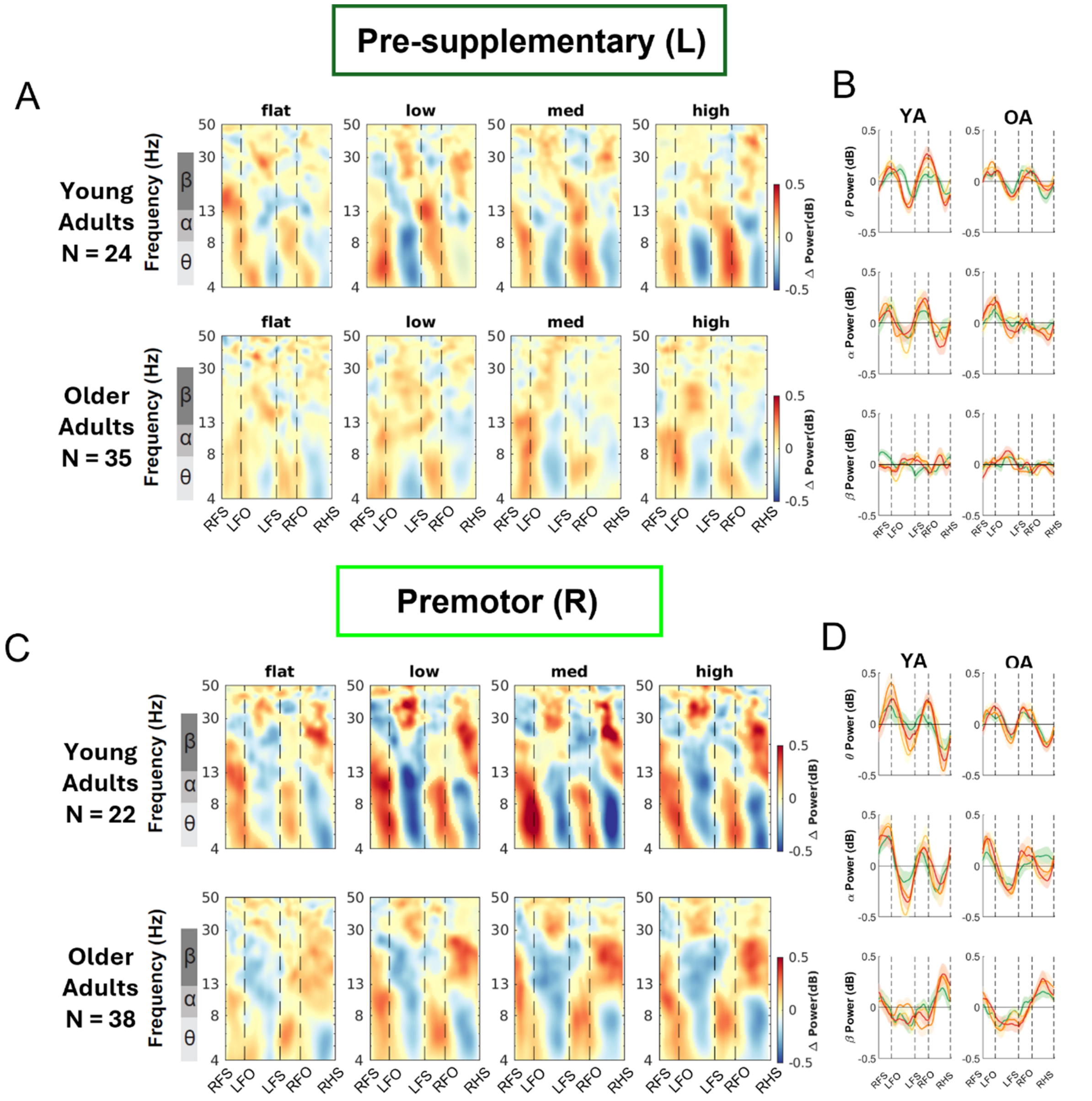


**Supplementary Figure 7**: Intra-stride event-related spectral perturbations (ERSPs) with respect to the average of condition at different terrains for young and older adults at (A) pre-supplementary and right premotor area (B), average intra-stride power fluctuations for each band at the (C) left pre-supplementary and right premotor area (D). (A, C) The ERSPs were displayed with the gait cycle on the x-axis (RFS: right foot strike; LTO: left toe off; LFO: left foot off; RFO: right foot off). All unmasked colors are statistically significant spectral power fluctuations relative to the mean power within the same condition. Colors indicate significant increases (red, synchronization) and decreases (blue, desynchronization) in spectral power from the average spectrum for all gait cycles to visualize intra-stride changes in the spectrograms. These data are significance masked ($p < 0.05$) through nonparametric bootstrapping with multiple comparison corrections using false discovery rate. (B, D) Average spectral perturbations for each band (theta, alpha, beta) across the gait cycle for young and older adults. YA: young adults. OA: older adults.

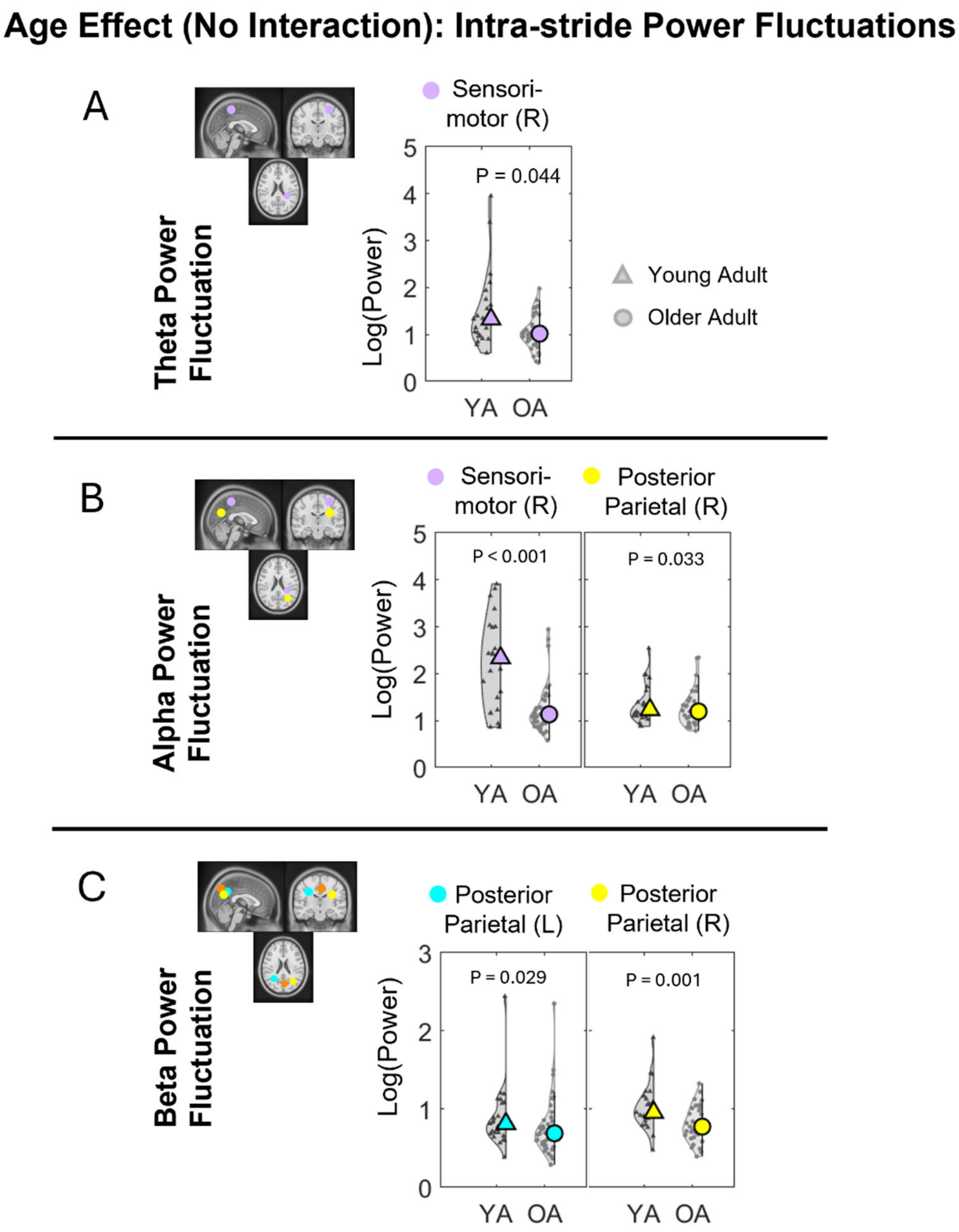


**Supplementary Figure 8**: Violin plots of the theta power fluctuations (A), alpha power fluctuations (B), and beta power fluctuations (C) for each age group after pooling all the terrain data for the corresponding brain clusters without an interaction effect. YA: younger adults, OA: older adults.

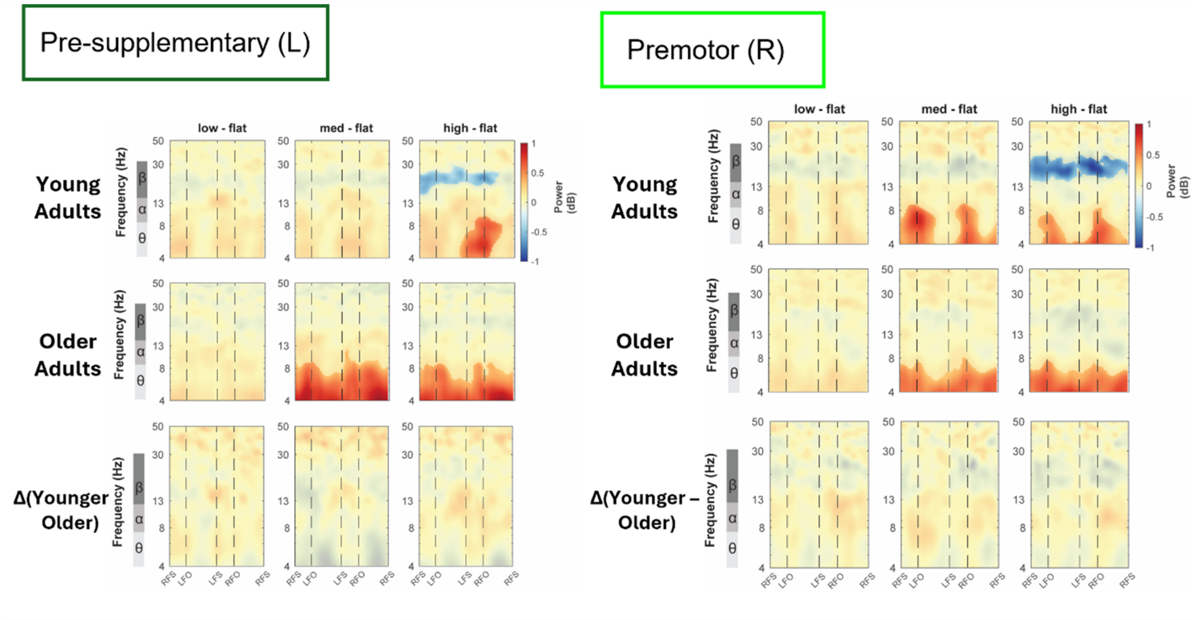


**Supplementary Figure 9**: Average event-related spectral perturbations (ERSP) with respect to the flat condition for each group and the difference between groups in the left pre-supplementary and right premotor. All unmasked colors are statistically significant spectral power fluctuations relative to the mean power within the same condition. The x-axes of the ERSPs are time in the gait cycle (RFS: right foot strike; LFO: left foot off; LFS: left foot strike; RFO: right foot off).

**Supplementary Table 1: Statistical table examining the effect of terrain, age, and the interaction on each behavioral outcome (*p<0.05, **p<0.01, ***p<0.001).**

| Measure | Effect | Fstats | dF | dF(res) | Pvalue | Sig | Partial eta$^2$ |
|---|---|---|---|---|---|---|---|
| Step Duration CoV | Terrain | 28.33 | 3 | 297 | <0.001 | *** | 0.22 |
| | Age | 0.01 | 1 | 166 | 0.920 | | 0 |
| | Treadmill Speed | 66.84 | 1 | 98 | <0.001 | *** | 0.41 |
| | Terrain:Age | 5.19 | 3 | 297 | 0.002 | ** | 0.05 |
| Anteroposterior Excursion CoV | Terrain | 88.14 | 3 | 300 | <0.001 | *** | 0.47 |
| | Age | 43.81 | 1 | 98 | <0.001 | *** | 0.31 |
| | Treadmill Speed | 25.54 | 1 | 98 | <0.001 | *** | 0.21 |
| Mediolateral Excursion CoV | Terrain | 10.47 | 3 | 297 | <0.001 | *** | 0.1 |
| | Age | 0.30 | 1 | 145.09 | 0.585 | | 0.002 |
| | Treadmill Speed | 10.80 | 1 | 98 | 0.001 | ** | 0.1 |
| | Terrain:Age | 3.99 | 3 | 297 | 0.008 | ** | 0.4 |

**Supplementary Table 2: Multiple comparison results for step duration CoV with significant interaction effect (*p<0.05, **p<0.01, ***p<0.001).**

| Measure | Group | Terrain | dF | t | pFDR- adjusted | Sig |
|---|---|---|---|---|---|---|
| Step Duration CoV | YA | Flat v. Low | 297 | -4.21 | <0.001 | *** |
| | | Flat v. Med | 297 | -6.44 | <0.001 | *** |
| | | Flat v. High | 297 | -8.85 | <0.001 | *** |
| | | Low v. Med | 297 | -2.23 | 0.027 | * |
| | | Low v. High | 297 | -4.64 | <0.001 | *** |
| | | Med v. High | 297 | -2.41 | 0.027 | * |
| | OA | Flat v. Low | 297 | -6.87 | <0.001 | *** |
| | | Flat v. Med | 297 | -11.16 | <0.001 | *** |
| | | Flat v. High | 297 | -19.66 | <0.001 | *** |
| | | Low v. Med | 297 | -4.29 | <0.001 | *** |
| | | Low v. High | 297 | -12.79 | <0.001 | *** |
| | | Med v. High | 297 | -8.50 | <0.001 | *** |

| Band | Terrain | Group | dF | t | pFDR- adjusted | Sig |
|---|---|---|---|---|---|---|
| Step Duration CoV | Flat | YA v. OA | 166 | 0.10 | 0.920 | |
| | Low | YA v. OA | 166 | -0.14 | 0.889 | |
| | Med | YA v. OA | 166 | -0.56 | 0.576 | |
| | High | YA v. OA | 166 | -2.74 | 0.007 | ** |

| Band | Group | Terrain | Z value | pFDR- adjusted | Sig |
|---|---|---|---|---|---|
| Step Duration CoV | YA v. OA | Low v. Flat | -0.30 | 0.765 | |
| | YA v. OA | Med v. Flat | -0.82 | 0.765 | |
| | YA v. OA | High v. Flat | -3.52 | 0.003 | ** |
| | YA v. OA | Med v. Low | -0.52 | 0.765 | |
| | YA v. OA | High v. Low | -3.22 | 0.006 | ** |
| | YA v. OA | High v. Med | -2.70 | 0.027 | * |

**Supplementary Table 3: Multiple comparison results for anteroposterior sacral excursion CoV with significant terrain effect (*p<0.05, **p<0.01, ***p<0.001).**

| Brain Area | Terrain | dF | t | pFDR- adjusted | Sig |
|---|---|---|---|---|---|
| Anteroposterior Excursion CoV | Flat v. Low | 300 | -7.467 | <0.001 | *** |
| | Flat v. Med | 300 | -10.839 | <0.001 | *** |
| | Flat v. High | 300 | -15.809 | <0.001 | *** |
| | Low v. Med | 300 | -3.371 | <0.001 | *** |
| | Low v. High | 300 | -8.342 | <0.001 | *** |
| | Med v. High | 300 | -4.971 | <0.001 | *** |

**Supplementary Table 4: Multiple comparison results for mediolateral sacral excursion CoV with significant interaction effect (*p<0.05, **p<0.01, ***p<0.001).**

| Measure | Group | Terrain | dF | t | pFDR- adjusted | Sig |
|---|---|---|---|---|---|---|
| Mediolateral Excursion CoV | YA | Flat v. Low | 297 | -2.21 | 0.084 | |
| | | Flat v. Med | 297 | -3.57 | 0.002 | ** |
| | | Flat v. High | 297 | -5.43 | <0.001 | *** |
| | | Low v. Med | 297 | -1.36 | 0.174 | |
| | | Low v. High | 297 | -3.22 | 0.006 | ** |
| | | Med v. High | 297 | -1.86 | 0.127 | |
| | OA | Flat v. Low | 297 | -6.77 | <0.001 | *** |
| | | Flat v. Med | 297 | -9.50 | <0.001 | *** |
| | | Flat v. High | 297 | -14.28 | <0.001 | *** |
| | | Low v. Med | 297 | -2.73 | 0.007 | ** |
| | | Low v. High | 297 | -7.52 | <0.001 | *** |
| | | Med v. High | 297 | -4.79 | <0.001 | *** |

| Band | Terrain | Group | dF | t | pFDR- adjusted | Sig |
|---|---|---|---|---|---|---|
| Mediolateral Excursion CoV | Flat | YA v. OA | 145 | 0.55 | 0.585 | |
| | Low | YA v. OA | 145 | -0.79 | 0.428 | |
| | Med | YA v. OA | 145 | -1.06 | 0.290 | |
| | High | YA v. OA | 145 | -1.84 | 0.068 | |

| Band | Group | Terrain | Z value | pFDR- adjusted | Sig |
|---|---|---|---|---|---|
| Mediolateral Excursion CoV | YA v. OA | Low v. Flat | -1.91 | 0.224 | |
| | YA v. OA | Med v. Flat | -2.29 | 0.110 | |
| | YA v. OA | High v. Flat | -3.39 | 0.004 | ** |
| | YA v. OA | Med v. Low | -0.38 | 0.704 | |
| | YA v. OA | High v. Low | -1.48 | 0.415 | |
| | YA v. OA | High v. Med | -1.10 | 0.541 | |

**Supplementary Table 5: Full statistical table for power spectral densities at theta, alpha, beta band at each brain area.** (SMA: sensorimotor; PP: posterior parietal; MCC: mid-cingulate; Pre-SMA: pre-supplementary motor)

| Brain Area | Band | Effect | Fstats | dF | dF(res) | Pvalue | Sig | Partial eta$^2$ |
|---|---|---|---|---|---|---|---|---|
| SMA L | Theta | Terrain | 0.94 | 3 | 225 | 0.421 | | 0.012 |
| | | Age | 5.25 | 1 | 73 | 0.025 | * | 0.067 |
| | | Treadmill Speed | 0.46 | 1 | 73 | 0.502 | | 0.006 |
| | Alpha | Terrain | 18.31 | 3 | 225 | < 0.001 | *** | 0.196 |
| | | Age | 0.28 | 1 | 73 | 0.598 | | 0.004 |
| | | Treadmill Speed | 0.61 | 1 | 73 | 0.438 | | 0.008 |
| | Beta | Terrain | 27.42 | 3 | 225 | < 0.001 | *** | 0.268 |
| | | Age | 1.15 | 1 | 73 | 0.288 | | 0.015 |
| | | Treadmill Speed | 1.79 | 1 | 73 | 0.185 | | 0.024 |
| SMA R | Theta | Terrain | 0.79 | 3 | 189 | 0.5 | | 0.012 |
| | | Age | 5.85 | 1 | 61 | 0.019 | * | 0.087 |
| | | Treadmill Speed | 1.91 | 1 | 61 | 0.172 | | 0.03 |
| | Alpha | Terrain | 18.85 | 3 | 189 | < 0.001 | *** | 0.23 |
| | | Age | 1.33 | 1 | 61 | 0.253 | | 0.021 |
| | | Treadmill Speed | 0.06 | 1 | 61 | 0.8 | | 0.001 |
| | Beta | Terrain | 19.14 | 3 | 189 | < 0.001 | *** | 0.233 |
| | | Age | 0.37 | 1 | 61 | 0.545 | | 0.006 |
| | | Treadmill Speed | 0.03 | 1 | 61 | 0.864 | | 0 |
| PP L | Theta | Terrain | 5.86 | 3 | 207 | 0.001 | *** | 0.078 |
| | | Age | 0.24 | 1 | 67 | 0.623 | | 0.004 |
| | | Treadmill Speed | 0.65 | 1 | 67 | 0.424 | | 0.01 |
| | Alpha | Terrain | 29.66 | 3 | 204 | < 0.001 | *** | 0.304 |
| | | Age | 0.36 | 1 | 67 | 0.553 | | 0.005 |
| | | Treadmill Speed | 0.69 | 1 | 67 | 0.409 | | 0.01 |
| | | Terrain:Age | 9.26 | 3 | 204 | < 0.001 | *** | 0.12 |
| | Beta | Terrain | 37.37 | 3 | 207 | < 0.001 | *** | 0.351 |
| | | Age | 0.29 | 1 | 67 | 0.592 | | 0.004 |
| | | Treadmill Speed | 2.01 | 1 | 67 | 0.161 | | 0.029 |
| PP R | Theta | Terrain | 3.97 | 3 | 171 | 0.009 | ** | 0.065 |
| | | Age | 0.05 | 1 | 55 | 0.829 | | 0.001 |
| | | Treadmill Speed | 1.56 | 1 | 55 | 0.218 | | 0.028 |
| | Alpha | Terrain | 18.32 | 3 | 168 | < 0.001 | *** | 0.247 |

| | | | | | | | | |
|---|---|---|---|---|---|---|---|---|
| | | Age | 0.22 | 1 | 55 | 0.642 | | 0.004 |
| | | Treadmill Speed | 0.09 | 1 | 55 | 0.759 | | 0.002 |
| | | Terrain:Age | 3.63 | 3 | 168 | 0.014 | * | 0.061 |
| | Beta | Terrain | 43.32 | 3 | 168 | < 0.001 | *** | 0.436 |
| | | Age | 1.61 | 1 | 55 | 0.209 | | 0.029 |
| | | Treadmill Speed | 0.06 | 1 | 55 | 0.809 | | 0.001 |
| | | Terrain:Age | 3.38 | 3 | 168 | 0.02 | * | 0.057 |
| MCC | Theta | Terrain | 3.59 | 3 | 177 | 0.015 | * | 0.057 |
| | | Age | 5.38 | 1 | 57 | 0.024 | * | 0.086 |
| | | Treadmill Speed | 0.59 | 1 | 57 | 0.445 | | 0.01 |
| | Alpha | Terrain | 4.16 | 3 | 177 | 0.007 | ** | 0.066 |
| | | Age | 0.11 | 1 | 57 | 0.744 | | 0.002 |
| | | Treadmill Speed | 0.06 | 1 | 57 | 0.805 | | 0.001 |
| | Beta | Terrain | 20.81 | 3 | 177 | < 0.001 | *** | 0.261 |
| | | Age | 4.71 | 1 | 57 | 0.034 | * | 0.076 |
| | | Treadmill Speed | 0.79 | 1 | 57 | 0.379 | | 0.014 |
| Occipital | Theta | Terrain | 29.46 | 3 | 225 | < 0.001 | *** | 0.282 |
| | | Age | 1.1 | 1 | 73 | 0.298 | | 0.015 |
| | | Treadmill Speed | 2.19 | 1 | 73 | 0.143 | | 0.029 |
| | Alpha | Terrain | 2.84 | 3 | 222 | 0.039 | * | 0.037 |
| | | Age | 0.71 | 1 | 73 | 0.402 | | 0.01 |
| | | Treadmill Speed | 0.12 | 1 | 73 | 0.729 | | 0.002 |
| | | Terrain:Age | 5.86 | 3 | 222 | 0.001 | *** | 0.073 |
| | Beta | Terrain | 47.75 | 3 | 222 | < 0.001 | *** | 0.392 |
| | | Age | 0.37 | 1 | 73 | 0.548 | | 0.005 |
| | | Treadmill Speed | 0.33 | 1 | 73 | 0.569 | | 0.004 |
| | | Terrain:Age | 6.36 | 3 | 222 | < 0.001 | *** | 0.079 |
| Pre-SMA | Theta | Terrain | 0.27 | 3 | 174 | 0.845 | | 0.005 |
| | | Age | 0.68 | 1 | 56 | 0.414 | | 0.012 |
| | | Treadmill Speed | 0 | 1 | 56 | 0.98 | | 0 |
| | Alpha | Terrain | 0.88 | 3 | 171 | 0.454 | | 0.015 |
| | | Age | 0.6 | 1 | 56 | 0.44 | | 0.011 |
| | | Treadmill Speed | 1.26 | 1 | 56 | 0.267 | | 0.022 |
| | | Terrain:Age | 2.83 | 3 | 171 | 0.04 | * | 0.047 |
| | Beta | Terrain | 8.64 | 3 | 174 | < 0.001 | *** | 0.13 |
| | | Age | 0.33 | 1 | 56 | 0.566 | | 0.006 |

| | | | | | | | | |
|---|---|---|---|---|---|---|---|---|
| | | Treadmill Speed | 0.7 | 1 | 56 | 0.405 | | 0.012 |
| Premotor | Theta | Terrain | 1.03 | 3 | 177 | 0.379 | | 0.017 |
| | | Age | 1.76 | 1 | 57 | 0.19 | | 0.03 |
| | | Treadmill Speed | 0.07 | 1 | 57 | 0.79 | | 0.001 |
| | Alpha | Terrain | 5.7 | 3 | 177 | 0.001 | *** | 0.088 |
| | | Age | 0.34 | 1 | 57 | 0.563 | | 0.006 |
| | | Treadmill Speed | 0.01 | 1 | 57 | 0.933 | | 0 |
| | Beta | Terrain | 7.86 | 3 | 177 | < 0.001 | *** | 0.118 |
| | | Age | 0.03 | 1 | 57 | 0.874 | | 0 |
| | | Treadmill Speed | 4.54 | 1 | 57 | 0.038 | * | 0.074 |

**Supplementary Table 6: Multiple comparisons between terrain conditions for each brain cluster that shows a significant terrain effect (*p<0.05, **p<0.01, ***p<0.001).**

| Band | Brain Area | Terrain | dF | t | pFDR- adjusted | Sig |
|---|---|---|---|---|---|---|
| Theta | PP L | Flat v. Low | 207 | 0.73 | 0.524 | |
| | | Flat v. Med | 207 | -0.64 | 0.524 | |
| | | Flat v. High | 207 | -3.21 | 0.008 | ** |
| | | Low v. Med | 207 | -1.37 | 0.519 | |
| | | Low v. High | 207 | -3.94 | 0.001 | *** |
| | | Med v. High | 207 | -2.57 | 0.044 | * |
| | PP R | Flat v. Low | 171 | -0.76 | 0.450 | |
| | | Flat v. Med | 171 | -1.53 | 0.384 | |
| | | Flat v. High | 171 | -3.29 | 0.007 | ** |
| | | Low v. Med | 171 | -0.77 | 0.450 | |
| | | Low v. High | 171 | -2.53 | 0.061 | |
| | | Med v. High | 171 | -1.76 | 0.321 | |
| | MCC | Flat v. Low | 177 | -0.84 | 0.785 | |
| | | Flat v. Med | 177 | -2.52 | 0.063 | |
| | | Flat v. High | 177 | -2.79 | 0.035 | * |
| | | Low v. Med | 177 | -1.68 | 0.286 | |
| | | Low v. High | 177 | -1.95 | 0.211 | |
| | | Med v. High | 177 | -0.27 | 0.785 | |
| | Occipital | Flat v. Low | 225 | -3.84 | <0.001 | *** |
| | | Flat v. Med | 225 | -6.92 | <0.001 | *** |
| | | Flat v. High | 225 | -8.77 | <0.001 | *** |
| | | Low v. Med | 225 | -3.07 | 0.005 | ** |
| | | Low v. High | 225 | -4.93 | <0.001 | *** |
| | | Med v. High | 225 | -1.86 | 0.065 | |
| **Band** | **Brain Area** | **Terrain** | **dF** | **t** | **pFDR- adjusted** | **Sig** |
| Alpha | SMA L | Flat v. Low | 225 | 4.18 | <0.001 | *** |
| | | Flat v. Med | 225 | 5.17 | <0.001 | *** |
| | | Flat v. High | 225 | 7.18 | <0.001 | *** |
| | | Low v. Med | 225 | 1.00 | 0.321 | |
| | | Low v. High | 225 | 3.01 | 0.009 | ** |
| | | Med v. High | 225 | 2.01 | 0.091 | |
| | SMA R | Flat v. Low | 189 | 3.70 | 0.001 | ** |
| | | Flat v. Med | 189 | 5.37 | <0.001 | *** |
| | | Flat v. High | 189 | 7.22 | <0.001 | *** |
| | | Low v. Med | 189 | 1.66 | 0.098 | |
| | | Low v. High | 189 | 3.51 | 0.002 | ** |
| | | Med v. High | 189 | 1.85 | 0.098 | |

| | | | | | | |
|---|---|---|---|---|---|---|
| | MCC | Flat v. Low | 177 | 2.52 | 0.063 | |
| | | Flat v. Med | 177 | 1.82 | 0.285 | |
| | | Flat v. High | 177 | 3.40 | 0.005 | ** |
| | | Low v. Med | 177 | -0.70 | 0.483 | |
| | | Low v. High | 177 | 0.88 | 0.483 | |
| | | Med v. High | 177 | 1.58 | 0.346 | |
| | Premotor | Flat v. Low | 177 | 2.66 | 0.034 | * |
| | | Flat v. Med | 177 | 2.88 | 0.022 | * |
| | | Flat v. High | 177 | 3.98 | 0.001 | *** |
| | | Low v. Med | 177 | 0.22 | 0.829 | |
| | | Low v. High | 177 | 1.31 | 0.548 | |
| | | Med v. High | 177 | 1.10 | 0.548 | |
| **Band** | **Brain Area** | **Terrain** | **dF** | **t** | **pFDR-** adjusted | **Sig** |
| Beta | SMA L | Flat v. Low | 225 | 4.31 | <0.001 | *** |
| | | Flat v. Med | 225 | 6.38 | <0.001 | *** |
| | | Flat v. High | 225 | 8.72 | <0.001 | *** |
| | | Low v. Med | 225 | 2.07 | 0.040 | * |
| | | Low v. High | 225 | 4.42 | <0.001 | *** |
| | | Med v. High | 225 | 2.35 | 0.040 | * |
| | SMA R | Flat v. Low | 189 | 3.70 | 0.001 | *** |
| | | Flat v. Med | 189 | 5.08 | <0.001 | *** |
| | | Flat v. High | 189 | 7.39 | <0.001 | *** |
| | | Low v. Med | 189 | 1.38 | 0.169 | |
| | | Low v. High | 189 | 3.69 | 0.001 | *** |
| | | Med v. High | 189 | 2.31 | 0.044 | * |
| | PP L | Flat v. Low | 207 | 4.86 | <0.001 | *** |
| | | Flat v. Med | 207 | 8.04 | <0.001 | *** |
| | | Flat v. High | 207 | 9.87 | <0.001 | *** |
| | | Low v. Med | 207 | 3.18 | 0.003 | ** |
| | | Low v. High | 207 | 5.01 | <0.001 | *** |
| | | Med v. High | 207 | 1.83 | 0.069 | |
| | MCC | Flat v. Low | 177 | 3.21 | 0.003 | ** |
| | | Flat v. Med | 177 | 3.27 | 0.003 | ** |
| | | Flat v. High | 177 | 7.84 | <0.001 | *** |
| | | Low v. Med | 177 | 0.06 | 0.953 | |
| | | Low v. High | 177 | 4.63 | <0.001 | *** |
| | | Med v. High | 177 | 4.57 | <0.001 | *** |
| | Pre-SMA | Flat v. Low | 174 | 1.91 | 0.162 | |
| | | Flat v. Med | 174 | 3.31 | 0.006 | ** |
| | | Flat v. High | 174 | 4.89 | <0.001 | *** |

| | | | | | | |
|---|---|---|---|---|---|---|
| | | Low v. Med | 174 | 1.40 | 0.162 | |
| | | Low v. High | 174 | 2.98 | 0.013 | * |
| | | Med v. High | 174 | 1.58 | 0.162 | |
| | Premotor | Flat v. Low | 177 | 2.33 | 0.071 | |
| | | Flat v. Med | 177 | 3.55 | 0.002 | ** |
| | | Flat v. High | 177 | 4.62 | <0.001 | *** |
| | | Low v. Med | 177 | 1.22 | 0.288 | |
| | | Low v. High | 177 | 2.28 | 0.071 | |
| | | Med v. High | 177 | 1.07 | 0.288 | |

**Supplementary Table 7: Multiple comparison results for left posterior parietal alpha band with significant interaction effect (*p<0.05, **p<0.01, ***p<0.001).**

| Band | Brain Area | Group | Terrain | dF | t | pFDR- adjusted | Sig |
|---|---|---|---|---|---|---|---|
| Alpha | PP L | YA | Flat v. Low | 204 | 4.92 | <0.001 | *** |
| | | | Flat v. Med | 204 | 8.21 | <0.001 | *** |
| | | | Flat v. High | 204 | 7.79 | <0.001 | *** |
| | | | Low v. Med | 204 | 3.29 | 0.004 | ** |
| | | | Low v. High | 204 | 2.87 | 0.009 | ** |
| | | | Med v. High | 204 | -0.43 | 0.671 | |
| | | OA | Flat v. Low | 204 | 1.61 | 0.217 | |
| | | | Flat v. Med | 204 | 2.18 | 0.121 | |
| | | | Flat v. High | 204 | 3.80 | 0.001 | ** |
| | | | Low v. Med | 204 | 0.57 | 0.570 | |
| | | | Low v. High | 204 | 2.19 | 0.121 | |
| | | | Med v. High | 204 | 1.62 | 0.217 | |

| Band | Brain Area | Terrain | Group | dF | t | pFDR- adjusted | Sig |
|---|---|---|---|---|---|---|---|
| Alpha | PP L | Flat | YA v. OA | 76.48 | 1.79 | 0.077 | |
| | | Low | YA v. OA | 76.48 | 0.60 | 0.548 | |
| | | Med | YA v. OA | 76.48 | -0.32 | 0.749 | |
| | | High | YA v. OA | 76.48 | 0.23 | 0.815 | |

| Band | Brain Area | Group | Terrain | Z value | pFDR- adjusted | Sig |
|---|---|---|---|---|---|---|
| Alpha | PP L | YA v. OA | Low v. Flat | -2.85 | 0.017 | * |
| | | YA v. OA | Med v. Flat | -5.08 | <0.001 | *** |
| | | YA v. OA | High v. Flat | -3.74 | <0.001 | *** |
| | | YA v. OA | Med v. Low | -2.23 | 0.078 | |
| | | YA v. OA | High v. Low | -0.89 | 0.374 | |
| | | YA v. OA | High v. Med | 1.34 | 0.362 | |

**Supplementary Table 8: Multiple comparison results for right posterior parietal alpha band with significant interaction effect (*p<0.05, **p<0.01, ***p<0.001).**

| Band | Brain Area | Group | Terrain | dF | t | pFDR- adjusted | Sig |
|---|---|---|---|---|---|---|---|
| Alpha | PP R | YA | Flat v. Low | 168 | 4.02 | <0.001 | *** |
| | | | Flat v. Med | 168 | 6.15 | <0.001 | *** |
| | | | Flat v. High | 168 | 6.13 | <0.001 | *** |
| | | | Low v. Med | 168 | 2.13 | 0.073 | |
| | | | Low v. High | 168 | 2.11 | 0.073 | |
| | | | Med v. High | 168 | -0.02 | 0.984 | |
| | | OA | Flat v. Low | 168 | 1.81 | 0.291 | |
| | | | Flat v. Med | 168 | 2.49 | 0.068 | |
| | | | Flat v. High | 168 | 2.95 | 0.022 | * |
| | | | Low v. Med | 168 | 0.69 | 0.648 | |
| | | | Low v. High | 168 | 1.15 | 0.648 | |
| | | | Med v. High | 168 | 0.46 | 0.648 | |

| Band | Brain Area | Terrain | Group | dF | t | pFDR- adjusted | Sig |
|---|---|---|---|---|---|---|---|
| Alpha | PP R | Flat | YA v. OA | 64.76 | 1.32 | 0.193 | |
| | | Low | YA v. OA | 64.76 | 0.47 | 0.643 | |
| | | Med | YA v. OA | 64.76 | -0.07 | 0.947 | |
| | | High | YA v. OA | 64.76 | 0.08 | 0.938 | |

| Band | Brain Area | Group | Terrain | Z value | pFDR- adjusted | Sig |
|---|---|---|---|---|---|---|
| Alpha | PP R | YA v. OA | Low v. Flat | -1.84 | 0.260 | |
| | | YA v. OA | Med v. Flat | -3.00 | 0.016 | * |
| | | YA v. OA | High v. Flat | -2.68 | 0.036 | * |
| | | YA v. OA | Med v. Low | -1.16 | 0.743 | |
| | | YA v. OA | High v. Low | -0.84 | 0.752 | |
| | | YA v. OA | High v. Med | 0.32 | 0.752 | |

**Supplementary Table 9: Multiple comparison results for occipital alpha band with significant interaction effect (*p<0.05, **p<0.01, ***p<0.001).**

| Band | Brain Area | Group | Terrain | dF | t | pFDR- adjusted | Sig |
|---|---|---|---|---|---|---|---|
| Alpha | Occipital | YA | Flat v. Low | 222 | 3.45 | 0.004 | ** |
| | | | Flat v. Med | 222 | 3.26 | 0.006 | ** |
| | | | Flat v. High | 222 | 3.20 | 0.006 | ** |
| | | | Low v. Med | 222 | -0.19 | 0.951 | |
| | | | Low v. High | 222 | -0.25 | 0.951 | |
| | | | Med v. High | 222 | -0.06 | 0.951 | |
| | | OA | Flat v. Low | 222 | 0.15 | 0.880 | |
| | | | Flat v. Med | 222 | -1.47 | 0.427 | |
| | | | Flat v. High | 222 | -1.93 | 0.276 | |
| | | | Low v. Med | 222 | -1.62 | 0.424 | |
| | | | Low v. High | 222 | -2.08 | 0.233 | |
| | | | Med v. High | 222 | -0.46 | 0.880 | |

| Band | Brain Area | Terrain | Group | dF | t | pFDR- adjusted | Sig |
|---|---|---|---|---|---|---|---|
| Alpha | Occipital | Flat | YA v. OA | 97.01 | 0.71 | 0.478 | |
| | | Low | YA v. OA | 97.01 | -0.92 | 0.360 | |
| | | Med | YA v. OA | 97.01 | -1.40 | 0.165 | |
| | | High | YA v. OA | 97.01 | -1.53 | 0.129 | |

| Band | Brain Area | Group | Terrain | Z value | pFDR- adjusted | Sig |
|---|---|---|---|---|---|---|
| Alpha | Occipital | YA v. OA | Low v. Flat | -2.71 | 0.027 | * |
| | | YA v. OA | Med v. Flat | -3.51 | 0.002 | ** |
| | | YA v. OA | High v. Flat | -3.72 | 0.001 | ** |
| | | YA v. OA | Med v. Low | -0.80 | 0.829 | |
| | | YA v. OA | High v. Low | -1.01 | 0.829 | |
| | | YA v. OA | High v. Med | -0.22 | 0.829 | |

**Supplementary Table 10: Multiple comparison results for left pre-supplementary motor band with significant interaction effect (*p<0.05, **p<0.01, ***p<0.001).**

| Band | Brain Area | Group | Terrain | dF | t | pFDR- adjusted | Sig |
|---|---|---|---|---|---|---|---|
| Alpha | Pre-SMA | YA | Flat v. Low | 171 | 0.38 | 0.707 | |
| | | | Flat v. Med | 171 | 1.21 | 0.707 | |
| | | | Flat v. High | 171 | -0.80 | 0.707 | |
| | | | Low v. Med | 171 | 0.83 | 0.707 | |
| | | | Low v. High | 171 | -1.18 | 0.707 | |
| | | | Med v. High | 171 | -2.01 | 0.279 | |
| | | OA | Flat v. Low | 171 | -0.54 | 0.587 | |
| | | | Flat v. Med | 171 | 0.65 | 0.587 | |
| | | | Flat v. High | 171 | 2.06 | 0.203 | |
| | | | Low v. Med | 171 | 1.20 | 0.587 | |
| | | | Low v. High | 171 | 2.61 | 0.060 | |
| | | | Med v. High | 171 | 1.41 | 0.587 | |

| Band | Brain Area | Terrain | Group | dF | t | pFDR- adjusted | Sig |
|---|---|---|---|---|---|---|---|
| Alpha | Pre-SMA | Flat | YA v. OA | 61.18 | -0.83 | 0.411 | |
| | | Low | YA v. OA | 61.18 | -1.04 | 0.300 | |
| | | Med | YA v. OA | 61.18 | -1.00 | 0.320 | |
| | | High | YA v. OA | 61.18 | -0.17 | 0.866 | |

| Band | Brain Area | Group | Terrain | Z value | pFDR- adjusted | Sig |
|---|---|---|---|---|---|---|
| Alpha | Pre-SMA | YA v. OA | Low v. Flat | -0.64 | 0.901 | |
| | | YA v. OA | Med v. Flat | -0.51 | 0.901 | |
| | | YA v. OA | High v. Flat | 1.93 | 0.214 | |
| | | YA v. OA | Med v. Low | 0.12 | 0.901 | |
| | | YA v. OA | High v. Low | 2.57 | 0.061 | |
| | | YA v. OA | High v. Med | 2.44 | 0.073 | |

**Supplementary Table 11: Multiple comparison results for right posterior parietal beta band with significant interaction effect (*p<0.05, **p<0.01, ***p<0.001).**

| Band | Brain Area | Group | Terrain | dF | t | pFDR- adjusted | Sig |
|---|---|---|---|---|---|---|---|
| Beta | PP R | YA | Flat v. Low | 168 | 2.83 | 0.011 | * |
| | | | Flat v. Med | 168 | 7.17 | <0.001 | *** |
| | | | Flat v. High | 168 | 8.11 | <0.001 | *** |
| | | | Low v. Med | 168 | 4.34 | <0.001 | *** |
| | | | Low v. High | 168 | 5.28 | <0.001 | *** |
| | | | Med v. High | 168 | 0.94 | 0.348 | |
| | | OA | Flat v. Low | 168 | 4.27 | <0.001 | *** |
| | | | Flat v. Med | 168 | 5.33 | <0.001 | *** |
| | | | Flat v. High | 168 | 6.54 | <0.001 | *** |
| | | | Low v. Med | 168 | 1.06 | 0.291 | |
| | | | Low v. High | 168 | 2.26 | 0.075 | |
| | | | Med v. High | 168 | 1.20 | 0.291 | |

| Band | Brain Area | Terrain | Group | dF | t | pFDR- adjusted | Sig |
|---|---|---|---|---|---|---|---|
| Beta | PP R | Flat | YA v. OA | 61.61 | 1.53 | 0.131 | |
| | | Low | YA v. OA | 61.61 | 1.79 | 0.079 | |
| | | Med | YA v. OA | 61.61 | 0.79 | 0.430 | |
| | | High | YA v. OA | 61.61 | 0.83 | 0.412 | |

| Band | Brain Area | Group | Terrain | Z value | pFDR- adjusted | Sig |
|---|---|---|---|---|---|---|
| Beta | PP R | YA v. OA | Low v. Flat | 0.67 | 0.935 | |
| | | YA v. OA | Med v. Flat | -1.90 | 0.205 | |
| | | YA v. OA | High v. Flat | -1.82 | 0.205 | |
| | | YA v. OA | Med v. Low | -2.58 | 0.059 | |
| | | YA v. OA | High v. Low | -2.50 | 0.062 | |
| | | YA v. OA | High v. Med | 0.08 | 0.935 | |

**Supplementary Table 12: Multiple comparison results for occipital beta band with significant interaction effect (*p<0.05, **p<0.01, ***p<0.001).**

| Band | Brain Area | Group | Terrain | dF | t | pFDR- adjusted | Sig |
|---|---|---|---|---|---|---|---|
| Beta | Occipital | YA | Flat v. Low | 222 | 6.64 | <0.001 | *** |
| | | | Flat v. Med | 222 | 8.36 | <0.001 | *** |
| | | | Flat v. High | 222 | 8.77 | <0.001 | *** |
| | | | Low v. Med | 222 | 1.72 | 0.174 | |
| | | | Low v. High | 222 | 2.13 | 0.102 | |
| | | | Med v. High | 222 | 0.42 | 0.678 | |
| | | OA | Flat v. Low | 222 | 5.32 | <0.001 | *** |
| | | | Flat v. Med | 222 | 4.74 | <0.001 | *** |
| | | | Flat v. High | 222 | 6.20 | <0.001 | *** |
| | | | Low v. Med | 222 | -0.59 | 0.559 | |
| | | | Low v. High | 222 | 0.88 | 0.559 | |
| | | | Med v. High | 222 | 1.46 | 4.359 | |

| Band | Brain Area | Terrain | Group | dF | t | pFDR- adjusted | Sig |
|---|---|---|---|---|---|---|---|
| Beta | Occipital | Flat | YA v. OA | 87.35 | 1.75 | 0.083 | |
| | | Low | YA v. OA | 87.35 | 0.66 | 0.511 | |
| | | Med | YA v. OA | 87.35 | -0.18 | 0.859 | |
| | | High | YA v. OA | 87.35 | 0.07 | 0.943 | |

| Band | Brain Area | Group | Terrain | Z value | pFDR- adjusted | Sig |
|---|---|---|---|---|---|---|
| Beta | Occipital | YA v. OA | Low v. Flat | -2.27 | 0.093 | |
| | | YA v. OA | Med v. Flat | -4.01 | <0.001 | *** |
| | | YA v. OA | High v. Flat | -3.49 | 0.002 | ** |
| | | YA v. OA | Med v. Low | -1.74 | 0.247 | |
| | | YA v. OA | High v. Low | -1.22 | 0.446 | |
| | | YA v. OA | High v. Med | 0.52 | 0.604 | |

**Supplementary Table 13 Full statistical table for power fluctuations at theta, alpha, beta band at each brain area.** (SMA: sensorimotor; PP: posterior parietal; MCC: mid-cingulate; Pre-SMA: pre-supplementary motor)

| Brain Area | Band | Effect | Fstats | dF | dF(res) | Pvalue | Sig | Partial eta$^2$ |
|---|---|---|---|---|---|---|---|---|
| SMA L | theta | Terrain | 0.41 | 3 | 225 | 0.745 | | 0.005 |
| | | Age | 1.29 | 1 | 73 | 0.261 | | 0.017 |
| | | Treadmill Speed | 1.03 | 1 | 73 | 0.314 | | 0.014 |
| | alpha | Terrain | 1.59 | 3 | 225 | 0.192 | | 0.021 |
| | | Age | 2.79 | 1 | 73 | 0.099 | | 0.037 |
| | | Treadmill Speed | 0.34 | 1 | 73 | 0.563 | | 0.005 |
| | beta | Terrain | 5.78 | 3 | 225 | 0.001 | *** | 0.072 |
| | | Age | 1.34 | 1 | 73 | 0.251 | | 0.018 |
| | | Treadmill Speed | 2.22 | 1 | 73 | 0.141 | | 0.029 |
| SMA R | theta | Terrain | 0.25 | 3 | 188 | 0.86 | | 0.004 |
| | | Age | 4.24 | 1 | 61 | 0.044 | * | 0.065 |
| | | Treadmill Speed | 0.29 | 1 | 61 | 0.592 | | 0.005 |
| | alpha | Terrain | 1.68 | 3 | 189 | 0.173 | | 0.026 |
| | | Age | 16.2 | 1 | 61 | < 0.001 | *** | 0.21 |
| | | Treadmill Speed | 0.15 | 1 | 61 | 0.699 | | 0.002 |
| | beta | Terrain | 0.89 | 3 | 189 | 0.449 | | 0.014 |
| | | Age | 3.47 | 1 | 61 | 0.067 | | 0.054 |
| | | Treadmill Speed | 0.09 | 1 | 61 | 0.77 | | 0.001 |
| PP L | theta | Terrain | 0.31 | 3 | 206 | 0.82 | | 0.004 |
| | | Age | 0.55 | 1 | 67 | 0.459 | | 0.008 |
| | | Treadmill Speed | 0.16 | 1 | 67 | 0.694 | | 0.002 |
| | alpha | Terrain | 0.84 | 3 | 206 | 0.471 | | 0.012 |
| | | Age | 3.85 | 1 | 67 | 0.054 | | 0.054 |
| | | Treadmill Speed | 0.75 | 1 | 67 | 0.388 | | 0.011 |
| | beta | Terrain | 0.47 | 3 | 206 | 0.706 | | 0.007 |
| | | Age | 4.97 | 1 | 67 | 0.029 | * | 0.069 |
| | | Treadmill Speed | 4.77 | 1 | 67 | 0.033 | * | 0.066 |
| PP R | theta | Terrain | 0.34 | 3 | 171 | 0.795 | | 0.006 |
| | | Age | 0.12 | 1 | 55 | 0.735 | | 0.002 |
| | | Treadmill Speed | 2.52 | 1 | 55 | 0.118 | | 0.044 |
| | alpha | Terrain | 0.59 | 3 | 171 | 0.623 | | 0.01 |
| | | Age | 4.77 | 1 | 55 | 0.033 | * | 0.08 |
| | | Treadmill Speed | 5.81 | 1 | 55 | 0.019 | * | 0.096 |
| | beta | Terrain | 0.47 | 3 | 171 | 0.702 | | 0.008 |
| | | Age | 12.18 | 1 | 55 | 0.001 | *** | 0.181 |

| | | | | | | | | |
|---|---|---|---|---|---|---|---|---|
| | | Treadmill Speed | 2.39 | 1 | 55 | 0.128 | | 0.042 |
| MCC | theta | Terrain | 0.06 | 3 | 177 | 0.978 | | 0.001 |
| | | Age | 0.02 | 1 | 57 | 0.892 | | 0 |
| | | Treadmill Speed | 0.08 | 1 | 57 | 0.782 | | 0.001 |
| | alpha | Terrain | 0.82 | 3 | 177 | 0.487 | | 0.014 |
| | | Age | 0.65 | 1 | 57 | 0.423 | | 0.011 |
| | | Treadmill Speed | 0.04 | 1 | 57 | 0.852 | | 0.001 |
| | beta | Terrain | 0.95 | 3 | 176 | 0.419 | | 0.016 |
| | | Age | 0.03 | 1 | 57 | 0.864 | | 0.001 |
| | | Treadmill Speed | 0.2 | 1 | 57 | 0.656 | | 0.004 |
| Occipital | theta | Terrain | 1.99 | 3 | 221 | 0.117 | | 0.026 |
| | | Age | 1.65 | 1 | 73 | 0.203 | | 0.022 |
| | | Treadmill Speed | 1.47 | 1 | 73 | 0.229 | | 0.02 |
| | | Terrain:Age | 4.8 | 3 | 221 | 0.003 | ** | 0.061 |
| | alpha | Terrain | 0.82 | 3 | 224 | 0.485 | | 0.011 |
| | | Age | 3.29 | 1 | 73 | 0.074 | | 0.043 |
| | | Treadmill Speed | 2.52 | 1 | 73 | 0.117 | | 0.033 |
| | beta | Terrain | 1.52 | 3 | 221 | 0.209 | | 0.02 |
| | | Age | 13.36 | 1 | 73 | < 0.001 | *** | 0.155 |
| | | Treadmill Speed | 4.1 | 1 | 73 | 0.047 | * | 0.053 |
| | | Terrain:Age | 3.24 | 3 | 221 | 0.023 | * | 0.042 |
| Pre-SMA | theta | Terrain | 0.79 | 3 | 174 | 0.501 | | 0.013 |
| | | Age | 1.51 | 1 | 56 | 0.224 | | 0.026 |
| | | Treadmill Speed | 0.12 | 1 | 56 | 0.727 | | 0.002 |
| | alpha | Terrain | 0.04 | 3 | 174 | 0.99 | | 0.001 |
| | | Age | 1.38 | 1 | 56 | 0.245 | | 0.024 |
| | | Treadmill Speed | 6.39 | 1 | 56 | 0.014 | * | 0.102 |
| | beta | Terrain | 1.49 | 3 | 174 | 0.22 | | 0.025 |
| | | Age | 2.92 | 1 | 56 | 0.093 | | 0.05 |
| | | Treadmill Speed | 5.04 | 1 | 56 | 0.029 | * | 0.083 |
| Premotor | theta | Terrain | 0.64 | 3 | 177 | 0.588 | | 0.011 |
| | | Age | 0.48 | 1 | 57 | 0.489 | | 0.008 |
| | | Treadmill Speed | 0.41 | 1 | 57 | 0.525 | | 0.007 |
| | alpha | Terrain | 0.52 | 3 | 177 | 0.667 | | 0.009 |
| | | Age | 0.43 | 1 | 57 | 0.514 | | 0.008 |
| | | Treadmill Speed | 3.06 | 1 | 57 | 0.085 | | 0.051 |
| | beta | Terrain | 0.89 | 3 | 177 | 0.448 | | 0.015 |
| | | Age | 1.8 | 1 | 57 | 0.185 | | 0.031 |
| | | Treadmill Speed | 2.65 | 1 | 57 | 0.109 | | 0.044 |

**Supplementary Table 14: Multiple comparison results for occipital theta band power fluctuations with significant interaction effect (*p<0.05).**

| Band | Brain Area | Group | Terrain | dF | t | pFDR- adjusted | Sig |
|---|---|---|---|---|---|---|---|
| Theta | Occipital | YA | Flat v. Low | 221 | -2.50 | 0.053 | |
| | | | Flat v. Med | 221 | -3.15 | 0.009 | ** |
| | | | Flat v. High | 221 | -3.28 | 0.007 | ** |
| | | | Low v. Med | 221 | -0.65 | 0.891 | |
| | | | Low v. High | 221 | -0.79 | 0.891 | |
| | | | Med v. High | 221 | -0.14 | 0.891 | |
| | | OA | Flat v. Low | 221 | 0.84 | 0.713 | |
| | | | Flat v. Med | 221 | 1.64 | 0.618 | |
| | | | Flat v. High | 221 | 0.48 | 0.713 | |
| | | | Low v. Med | 221 | 0.80 | 0.713 | |
| | | | Low v. High | 221 | -0.37 | 0.713 | |
| | | | Med v. High | 221 | -1.17 | 0.713 | |

| Band | Brain Area | Terrain | Group | dF | t | pFDR- adjusted | Sig |
|---|---|---|---|---|---|---|---|
| Theta | Occipital | Flat | YA v. OA | 149 | -1.03 | 0.303 | |
| | | Low | YA v. OA | 149 | 1.31 | 0.192 | |
| | | Med | YA v. OA | 149 | 2.24 | 0.027 | * |
| | | High | YA v. OA | 149 | 1.71 | 0.090 | |

| Band | Brain Area | Group | Terrain | Z value | pFDR- adjusted | Sig |
|---|---|---|---|---|---|---|
| Theta | Occipital | YA v. OA | Low v. Flat | 2.52 | 0.047 | * |
| | | YA v. OA | Med v. Flat | 3.52 | 0.003 | ** |
| | | YA v. OA | High v. Flat | 2.94 | 0.016 | * |
| | | YA v. OA | Med v. Low | 1.00 | 0.672 | |
| | | YA v. OA | High v. Low | 0.42 | 0.672 | |
| | | YA v. OA | High v. Med | -0.58 | 0.672 | |

**Supplementary Table 15: Multiple comparison results for occipital beta band power fluctuations with significant interaction effect (*p<0.05, **p<0.01, ***p<0.001).**

| Band | Brain Area | Group | Terrain | dF | t | pFDR- adjusted | Sig |
|---|---|---|---|---|---|---|---|
| Beta | Occipital | YA | Low v. Flat | 221 | -1.61 | 0.437 | |
| | | | Med v. Flat | 221 | -2.32 | 0.106 | |
| | | | High v. Flat | 221 | -2.86 | 0.028 | * |
| | | | Med v. Low | 221 | -0.71 | 0.590 | |
| | | | High v. Low | 221 | -1.25 | 0.590 | |
| | | | High v. Med | 221 | -0.54 | 0.590 | |
| | | OA | Low v. Flat | 221 | 1.69 | 0.556 | |
| | | | Med v. Flat | 221 | 0.50 | 0.619 | |
| | | | High v. Flat | 221 | 1.03 | 0.619 | |
| | | | Med v. Low | 221 | -1.20 | 0.619 | |
| | | | High v. Low | 221 | -0.66 | 0.619 | |
| | | | High v. Med | 221 | 0.54 | 0.619 | |

| Band | Brain Area | Terrain | Group | dF | t | pFDR- adjusted | Sig |
|---|---|---|---|---|---|---|---|
| Beta | Occipital | Flat | YA v. OA | 155 | 1.20 | 0.234 | |
| | | Low | YA v. OA | 155 | 3.39 | 0.001 | *** |
| | | Med | YA v. OA | 155 | 3.28 | 0.001 | ** |
| | | High | YA v. OA | 155 | 3.99 | < 0.001 | *** |

| Band | Brain Area | Group | Terrain | Z value | pFDR- adjusted | Sig |
|---|---|---|---|---|---|---|
| Beta | Occipital | YA v. OA | Low v. Flat | 2.30 | 0.109 | |
| | | YA v. OA | Med v. Flat | 2.17 | 0.119 | |
| | | YA v. OA | High v. Flat | 2.92 | 0.021 | * |
| | | YA v. OA | Med v. Low | -0.12 | 0.902 | |
| | | YA v. OA | High v. Low | 0.63 | 0.902 | |
| | | YA v. OA | High v. Med | 0.75 | 0.902 | |
